# Histological validation of per-bundle water diffusion metrics within a region of fiber crossing following axonal degeneration

**DOI:** 10.1101/571539

**Authors:** Gilberto Rojas-Vite, Ricardo Coronado-Leija, Omar Narvaez-Delgado, Alonso Ramírez-Manzanares, José Luis Marroquín, Ramsés Noguez-Imm, Marcos L. Aranda, Benoit Scherrer, Jorge Larriva-Sahd, Luis Concha

**Author notes:** Corresponding author: *Email address:* (Luis Concha). Equal contributions (Gilberto Rojas-Vite, Ricardo Coronado-Leija).

## Abstract

Micro-architectural characteristics of white matter can be inferred through analysis of diffusion-weighted magnetic resonance imaging (dMRI). The diffusion-dependent signal can be analyzed through several methods, with the tensor model being the most frequently used due to its straightforward interpretation and relaxed acquisition parameters. While valuable information can be gained from the tensor-derived metrics in regions of homogeneous tissue organization, this model does not provide reliable microstructural information at crossing fiber regions, which are pervasive throughout human white matter. Several multiple fiber models have been proposed that seem to overcome the limitations of the tensor, with few providing per-bundle dMRI-derived metrics. However, biological interpretations of such metrics are limited by the lack of histological confirmation. To this end, we developed a straightforward biological validation framework. Unilateral retinal ischemia was induced in ten rats, which resulted in axonal (Wallerian) degeneration of the corresponding optic nerve, while the contralateral was left intact; the intact and injured axonal populations meet at the optic chiasm as they cross the midline, generating a fiber crossing region in which each population has different diffusion properties. Five rats served as controls. High-resolution *ex vivo* dMRI was acquired five weeks after experimental procedures. We correlated and compared histology to per-bundle descriptors derived from three novel methodologies for dMRI analysis (constrained spherical deconvolution and two multi-tensor representations). We found a tight correlation between axonal density (as evaluated through automatic segmentation of histological sections) with per-bundle apparent fiber density and fractional anisotropy (derived from dMRI). The multi-fiber methods explored were able to correctly identify the damaged fiber populations in a region of fiber crossings (chiasm). Our results provide validation of metrics that bring substantial and clinically useful information about white-matter tissue at crossing fiber regions. Our proposed validation framework is useful to validate other current and future dMRI methods.

## 1. Introduction

Magnetic resonance imaging (MRI) provides different contrast mechanisms that convey important information regarding tissue. Of particular interest in the quest for biomarkers is the ability to non-invasively infer the architectural and cellular characteristics of tissue, as a means for detection and long-term follow-up of disease-related abnormalities. Brain imaging has seen a host of quantitative methods to query tissue composition (Cercignani et al., 2018). The common denominator of many methods is their capacity to resolve information beyond their spatial resolution limit. Diffusion-weighted MRI (dMRI), in particular, has gained great traction for the study of cerebral white matter and, aided by tractography algorithms that enable the depiction of white matter bundles, has enabled the field of anatomical brain connectivity (Sporns, 2011; Dell’Acqua and Catani, 2012). While the study of trajectories of large white matter pathways provides valuable information that can be used in the clinic (e.g., for surgical planning), a wealth of information is contained within each image voxel (Concha, 2014; Paus, 2018; Dyrby et al., 2018).

The self-diffusion of water molecules in tissue is modulated by the barriers that molecules encounter over a period of time (Beaulieu, 2002). In tissues where the organization of cellular membranes and other components are organized in a very coherent fashion with evident textural orientation, water will not diffuse randomly, but tend to follow the architecture of tissue. Through many observations, dMRI queries the profile of water diffusion in three dimensions, and mathematical models are used to provide a compiled, often simplified, view of the dMRI signal at each voxel. The diffusion tensor (Basser et al., 1994) is a simple and intuitive model that has proven useful for the identification of features of white matter bundles that are modulated by disease, neurodevelopment, plasticity, and aging. Several studies have provided direct evidence that tensor metrics are related to histological characteristics in normal and affected white matter (Leergaard et al., 2010; Concha et al., 2010; Sierra et al., 2015; Khan et al., 2015; Salo et al., 2018). Diffusion is modeled through the water molecular displacement co-variance matrix, and represented as an ellipsoid oriented parallel to the main direction of the nervous tissue (axons). Thus, reductions of the largest matrix eigenvector (λ_1_=λ_║_) reflect axonal beading or fragmentation (Budde et al., 2007; Song et al., 2003; Liu et al., 2013), whereas increased perpendicular diffusivity (λ_⊥_ = (λ_2_ + λ_3_)/2) can reflect one or several mechanisms, including loss of axons, changes in average axonal diameter, loss of fiber coherence, fiber dispersion, or myelin loss (Song et al., 2003; Tyszka et al., 2006; Budde et al., 2007; Concha et al., 2010; Klawiter et al., 2011). This intuitive interpretation (in spite of its ambiguities) and low data acquisition requirements propel the tensor model to this day (O’Donnell and Westin, 2011).

One of the main assumptions made in the tensor model is the existence of a single dominant fiber population within any given voxel. This, however, is not the case in the majority of human white matter, which contains more than one fiber population with different configurations (Jeurissen et al., 2013). The tensor model cannot provide reliable information in these areas and is (at its best) not useful, or (at its worse) misleading (Tournier et al., 2011; Jones et al., 2013). For example, degeneration of one fiber population within a voxel occupied by two crossing populations leads to increased diffusion anisotropy (Douaud et al., 2011), typically and over-simplistically associated with healthy tissue (Jones et al., 2013).

Alternatives to the tensor model abound (Alexander, 2005; Tournier et al., 2011; Panagiotaki et al., 2012; Daducci et al., 2014). Some methods use diffusion metrics directly as biomarkers of tissue characteristics, whereas others attempt to build specific biophysical models for tissue parameters (Jespersen, 2018). In general, they aim to resolve the correct number and orientation of axonal populations and, ideally, provide per-bundle fiber characteristics to improve tissue microstructure estimation or tractogram construction for brain connectivity analyses (Dell’Acqua and Catani, 2012). It has been shown that advanced dMRI models provide information about bundle orientation in fiber crossing regions, which corresponds to quantitative histology (Leergaard et al., 2010; Budde and Annese, 2013). However, it is often crucial not only to correctly infer the orientation of fiber populations, but also to provide microstructural information of each fiber system in a crossing region. While numerical simulation studies have provided insight into the validity of multi-fiber dMRI reconstruction methods (Alexander, 2005), direct validations with ground-true methods (i.e., histology) are needed for appropriate biological interpretations. Recent methods have shown that they can estimate certain tissue characteristics, such as intracellular volume fractions and bundle dispersion (Zhang et al., 2012; Daducci et al., 2015). However, those methods estimate tissue parameters as an average over the whole voxel, and cannot disentangle characteristics for each axonal bundle if more than one occupies a particular voxel.

In this work we show the ability of a subset of multi–fiber reconstruction methods to capture histological features for individual fiber bundles in fiber crossing regions. By extending the well-known rodent model of unilateral retinal ischemia (Adachi et al., 1996), which has been extensively used to validate dMRI methods (Song et al., 2003; Sun et al., 2008), we show that dMRI can correctly identify and quantify a single population of degenerated axons from a population of intact axons within the optic chiasm.

## 2. Methods

### 2.1. Animal preparation

We used the retinal ischemia model to induce axonal (Wallerian) degeneration of axons emanating from the retina towards the brain (Adachi et al., 1996; Song et al., 2003). Ischemia was induced unilaterally, thus providing one injured and one intact nerve for each animal. Optic chiasm analysis allows for the evaluation of crossing fibers, as axons from each eye cross the midline at this level. The proportion of axons that cross the midline is much larger in rodents than in humans, being over 90% in the former, and roughly half in the latter (Jeffery and Erskine, 2005). Thus, unilateral retinal ischemia results in degeneration of practically half of the total number of axons within the chiasm (Fig. 1). Fifteen adult female Wistar rats were studied (age 16 to 18 weeks; weight: 354±59 g); axonal degeneration was induced in ten, and five served as controls. To cause unilateral retinal ischemia, animals were anesthetized with a ketamine/xylazine solution administered I.P. and then placed in a stereotaxic frame. A 32-gauge needle was inserted into the anterior chamber of the right eye of each rat, and connected to a reservoir with saline solution that was elevated until an in-line pressure monitor indicated 120 mmHg (higher than systolic pressure); this pressure was maintained for 90 minutes. Retinal pallor was evident throughout the experiment, confirming lack of retinal blood flow. After this period the cannula was carefully extracted and topical antibiotics and analgesics were administered. Animals were returned to their cages to recover with *ad libitum* access to food and water. Five weeks after treatment, animals were deeply anesthetized I.P. with ketamine/xylazine and intracardially perfused with 4% paraformaldehyde (PFA) and 2.5 % glutaraldehyde with Gadolinium (0.2 mM) (D’Arceuil et al., 2007). Brains were carefully extracted leaving the optic nerves and chiasm intact. To prevent the optic nerves from floating, their most distal portions were attached to the ventral aspect of the olfactory bulbs using cyanoacrylate, and specimens were placed in PFA at 4 °C until imaging. All procedures were performed in compliance with international ethics and animal care guidelines, and the study was approved by the Bioethics Committee of the Institute of Neurobiology, Universidad Nacional Autónoma de México (protocol 096.A).

**Figure 1:**
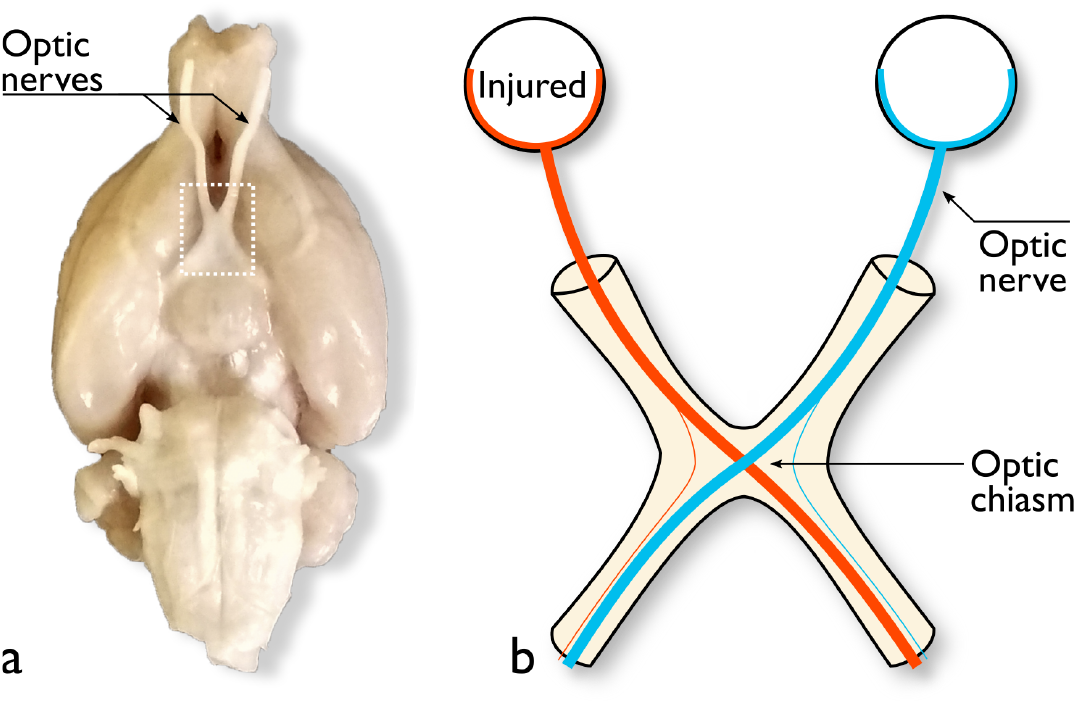
Biological model. a: The ventral aspect of a fixed rat brain shows the two optic nerves affixed to the basal aspect of the olfactory bulbs, and meeting at the midline to form the optic chiasm (dotted rectangle). b: Schematic representation of the retinal ischemia model, which induces Wallerian degeneration of axons traveling through one optic nerve (orange). The contralateral optic nerve remains intact (blue). At the level of the optic chiasm, the majority of axons cross the midline, thus providing an anatomical region of known fiber crossings where a single fiber population has been affected.

### 2.2. Imaging

Brains were scanned using a 7 T Bruker Pharmascan 70/16 with 760 mT/m maximum gradient amplitude using a combination of a 72 mm inner-diameter circularly polarized radio-frequency coil (for transmission), and a rat head 2×2 surface array rat coil (receive-only). The specimens were taken out of refrigeration two hours before imaging, placed in a plastic tube and immersed in perfluoropolyether (Fomblin Y, Sigma-Aldrich), oriented such that the chiasm and optic nerves were in close proximity to the receiving coil. A *B*_0_ map was obtained and used to aid automatic shimming before dMRI acquisition. Images were acquired with 125 × 125 × 125 μm^3^ resolution using a 3-dimensional echo-planar acquisition with 8 segments (TR/TE=250/21 ms; NEX=1). We acquired diffusion-weighted images (DWI) in 80 different directions, each with a *b* value of 2000 and 2500 s/mm^2^. (δ/Δ=3.1/10 ms), and 20 non-diffusion weighted images. Total scanning time was 15 hours, and scanning was performed at room temperature (21 °C) with fluctuations of less than 1.5 °C (measured twice with an MRI-compatible thermometer [Small Animal Instruments, Inc., Stony Brook, NY] placed inside the scanner bore). Imaging sessions were performed 24±21 (range 2-63) days after intracardiac perfusion, with specimens kept in PFA at 4 °C during this period. After scanning, specimens were returned to PFA and refrigerated until preparation for histology.

### 2.3. Analysis of dMRI

To compute dMRI-based microstructure estimators, we used three different modern approaches: constrained spherical deconvolution (CSD) (Tournier et al., 2011), multi-resolution discrete search (MRDS) (Coronado-Leija et al., 2017), and distribution of anisotropic microstructural environments (DIAMOND) (Scherrer et al., 2016). CSD is currently one of the most widely used multi–fiber methods, and uses a fixed response function to explain dMRI signals. The multi-tensor model solved by MRDS is the direct extension to the diffusion tensor, providing related per-bundle estimators. DIAMOND goes one step further by adding dispersion parameters and an isotropic compartment in its formulation. No prior information was provided to the methods about tissue orientation or characteristics. Model fitting was performed voxel-wise (i.e., no spatial smoothness or spatial coherence was imposed).

#### 2.3.1. Pre-processing

Images were first denoised (Veraart et al., 2016), then bias field inhomogeneities were corrected. The bias field was estimated using the N4 algorithm (Tustison et al., 2010) on the average *b* = 0 s/mm^2^ volume, and the resulting correction factor applied to all volumes. Finally, geometric distortions induced by eddy currents, as well as image drift during the long acquisition period, were corrected by registering each volume to the average *b* = 0 s/mm^2^ volume using a linear transformation with 12 degrees of freedom using tools available in the *fsl* suite (Jenkinson et al., 2002).

#### 2.3.2. CSD

Voxels with a single fiber population within a manually-defined region encompassing the intact optic nerve(s) were identified for each specimen using an automated approach (Tournier et al., 2013) (number of single-fiber voxels identified = 104.4 ± 40.8). The median *b* = 0 s/mm^2^ intensity of those voxels was used to normalize the dMRI signal. Next, the response function was estimated from the dMRI signal within the same voxels. Fiber orientation distribution (FOD) functions were obtained for white matter, gray matter and CSF using the method proposed by Jeurissen et al. (2014). The resulting response functions were averaged across all animals to obtain a single response function for each tissue type, which was used to estimate the FODs for each animal. In this work we focused on the FOD derived from white matter and report only the results using the corresponding response function. Voxel-wise FOD complexity was estimated (Riffert et al., 2014). Per-bundle (i.e., fixel-wise) metrics were derived by FOD segmentation (Smith et al., 2013), from which we calculated apparent fiber density (AFD), peak AFD, and dispersion (Raffelt et al., 2012; Dell’Acqua et al., 2013; Riffert et al., 2014)).

#### 2.3.3. MRDS

We computed parameters of the multi-tensor model at single-fiber and crossing-fiber regions using the multi-resolution discrete search (MRDS) fitting method (Coronado-Leija et al., 2017). This state-of-the-art fitting technique provides robust estimates of specific intra-voxel diffusion descriptors, per-bundle volume fractions (*α*), orientations, and axial and radial diffusivities (λ_║_ and λ_⊥_, respectively). The recovered diffusion information allows, in principle, to detect diffusion abnormalities of individual fiber populations. The initial response function (a radially symmetric tensor shape) was computed by averaging the single-tensor eigenvalues of voxels that likely contain a single bundle. Single-fiber voxels were identified by fitting the single tensor model to all voxels in a region of interest (ROI) delineated at the intact optic nerve(s) for each specimen, from which we computed fractional anisotropy (FA), and the average and standard deviation of the mean diffusivity (*μ_MD_* and *σ_MD_*, respectively), as well as the linear planar and spherical coefficients (λ_1_, λ_*p*_ λ_*s*_, respectively) (Westin et al., 2002). The single bundle voxels were defined as those having mean diffusivity (MD) in the range [*μ_MD_* − *σ_MD_*,*μ_MD_* + *σ_MD_*], FA≥0.7, λ_*l*_ > λ_*p*_ and λ_*l*_ > λ_*s*_. Later, the MRDS updated the per-bundle tensor response functions (the individual eigenvalues, and principal diffusion directions) associated with N=1,2 and 3 tensors per voxel. The final result is computed by a model selection stage that used the *ad hoc* statistical F-test for nested models (see Coronado-Leija et al. (2017) for further details). For each bundle, we report FA, MD, λ_║_, λ_⊥_, and compartment size (*α*).

#### 2.3.4. DIAMOND

We evaluated the DIAMOND (distribution of anisotropic microstructural environment in diffusion imaging) approach which models the diffusion arising from each fiber using a continuous, peak-shaped, statistical distribution of diffusion tensors (Scherrer et al., 2016). This allows modeling of both the average fiber 3-D diffusivity and its microstructural heterogeneity, and accounts for the non-monoexponential decay in tissues. DIAMOND has been shown to be a good predictor of the diffusion signal (Ferizi et al., 2017) and can be seen as an extension of the multi-tensor model, which is a DIAMOND model with infinitely concentrated tensor distributions representing purely homogeneous compartments. DIAMOND is able to capture variations around the average diffusivity and is expected to better estimate the average diffusivity itself.

Model selection to determine the number of fibers at each voxel was achieved using the Akaike criterion (Akaike, 1974). The statistical distribution of tensors was chosen to be asymmetric to model decoupled axial and radial heterogeneity (Scherrer et al., 2017), with a freely-diffusing compartment with a fixed *D_iso_* = 2.0668 × 10^−3^mm^2^/s, for which its isotropic fraction *f_iso_* was estimated. Bundle-specific diffusivities were calculated by estimating the eigenvalues of the expectation of each distribution which is a tensor that represents the average compartment-specific diffusivity. Metrics were derived for each fiber bundle, namely FA, MD, λ_║_, λ_⊥_, *α*, and heterogeneity indices for λ_║_ and λ_⊥_ (λ_║*HEI*_ and λ_⊥_, respectively).

### 2.4. Histology

#### 2.4.1. Tissue preparation

Following dMRI acquisition, the specimens were returned to PFA until processing (time between dMRI and tissue preparation was 47±29 [range 11-118] days). The optic nerves and chiasm were separated from the brain to proceed with histological preparation. Samples were stained with Osmium tetraoxide, then washed with phosphate buffered saline (0.1 M) and dehydrated with acetone in a gradient of concentrations (60,70,80 and 90%) until absolute acetone. After, tissue was embedded in a 2:1 epoxy resin/acetone solution for approximately 18 hours. For polymerization, samples were placed in a mold with epoxy resin and kept at 60 °C. Optic nerves were sectioned (500 nm thick) perpendicular to their long axis, whereas the optic chiasm was sagittally sectioned at the midline. Each section was stained with a solution of toluidine blue and sodium borate (both 0.5%) to continue with observation and photomicrographic acquisition using an optical microscope. For illustrative purposes, selected specimens underwent electron microscopy, using silver sections at a nominal thickness of 70 nm; these sections were imaged with a JEOL 1010 electron microscope operated at 80 kv.

#### 2.4.2. Quantitative evaluation

After staining optic nerve sections, we took photomicrograph tiles using a vertical Zeiss Axio Imager microscope with a motorized stage at 63× magnification. Illumination, contrast, focus and exposure were adjusted for every frame. Tiles were stitched with the “Grid/collection stitching” plugin (Preibisch et al., 2009) available in Fiji software (Schindelin et al., 2012) to obtain three mosaics of 3886× 3848 pixels of complete sections for every optic nerve. Quantitative evaluation of the mosaics was performed with AxonSeg (Zaimi et al., 2016), using a single set of segmentation parameters for both control and experimental conditions. We obtained total axon count, axon diameter, and axon+myelin diameter. From these metrics and total optic nerve area, we also obtained axon density, axon volume fraction (AVF), myelin thickness, g-ratio, and myelin volume fraction (MVF). The complicated architectural disposition of axons in the optic chiasm, and the difficulty to obtain sections oriented orthogonally to both axon bundles, precluded direct quantitative evaluation of this region. We could not find a single set of parameters for automatic segmentation of axons that produced reliable estimations for the two axonal populations simultaneously, particularly in the injured condition. To overcome this problem, and given that in rat brains nearly all axons from each optic nerve cross the midline (Jeffery and Erskine, 2005), estimations of the two axonal populations in the chiasm were done indirectly, by referring to the corresponding optic nerve quantifications.

### 2.5. Data availability

All raw dMRI, as well as all photomicrographs and corresponding automatic segmentations, are available in the White Matter Microscopy Database (https://osf.io/yp4qg/) (Cohen-Adad et al., 2017)

### 2.6. Statistical analyses

Manually-drawn ROIs were delineated at the level of each optic nerve and at the center of the chiasm. Optic nerve ROIs were 229±103 voxels in size, whereas chiasm ROIs had 27±2 voxels. For each ROI, we calculated the mean and standard deviation of each of the metrics provided by the three multi-fiber dMRI reconstruction methods. For voxels in optic nerves with more than one fiber bundle (identified more frequently in injured nerves), we used only the metrics for the bundle with the largest AFD (CSD) and α (MRDS and DIAMOND) in the computations. For voxels in the chiasm, the metrics for the obtained bundles were identified as *intact* and *injured* based on their principal diffusion orientation with respect to the corresponding optic nerve. To perform this clustering, we used orientations of bundles with the largest AFD and α (up to three) in the ROI to compute the centroid of orientations of the bundles using the k-means method (*k* = 2). These centroids were then used to discriminate between intact bundle metrics and injured bundle metrics in each voxel. Finally, the mean and standard deviation of diffusion metrics were computed for each bundle.

Given the histology-based values and the metrics obtained from the dMRI multi-fiber methods, the Pearson correlation coefficient was computed for each histology/dMRI pair. In nerves, this computation was straightforward: the histology-based values in one nerve were assigned to the dMRI descriptor values corresponding to the same nerve. This was done for all rats, giving a total of 30 points (15 rats × 2 nerves) that were used to calculate the Pearson correlation coefficient. The corresponding p-values were computed using permutation testing with 10,000 permutations to create the null hypothesis distribution. For the chiasm, a similar procedure was used, but because no quantitative histological information was obtained for the chiasm, the dMRI metric values for each crossing bundles were assigned to the histology-based values for the corresponding nerve. For voxel-wise metrics, where no bundle-specific information exists, the dMRI descriptors were correlated to the mean of the histology-based values of both nerves.

#### 2.6.1. Bootstrap aggregating for sensitivity evaluation

To study the sensitivity of the provided dMRI descriptors to the reduction of volumes in the data, as well as to characterize the statistical performance of the dMRI models when exposed to a large set of experiments, we generated subsets of the data with replacement using the Bootstrap Aggregating Method (BAM; also known as Bagging) (Breiman, 1996). BAM allows the estimation of the average quality of the dMRI methodologies for a large number of experiments when only a few samples are available. We created 100 samples of reduced dMRI data for five cases, each one with 30, 40, 50, 60 or 70 volumes per shell, for which the corresponding gradients are evenly distributed in the half sphere. CSD, MRDS and DIAMOND models were fitted to these data subsets and the statistical analyses described above were performed with the resulting dMRI descriptors. By simulating a large collection of data sets with the BAM procedure we were able to evaluate the quality of the results in clinically-viable settings (i.e., for a reduced number of dMRI measurements).

## 3. Results

Injured optic nerves showed marked atrophy when compared to the intact optic nerves (Figs. 2a and e). Evaluation of intact optic nerves using light microscopy revealed that the distribution of axons throughout the optic nerve is homogeneous with fibers grouped into coalescent fascicles bounded by glial processes conveying capillary blood vessels (Fig. 2b). Automatic segmentation of axons in the intact nerve shows that most axons have diameters in the range of one micron, with an average myelin thickness of 0.34 *μ*m (Figs. 2, 4, and Supplementary Fig. 4). Injured optic nerves showed an abnormal microstructure (Fig. 2f) involving axons and surrounding glial elements. High magnification views (Fig. 2f) show a marked reduction of axonal density and the presence of bizarre, dense, retracted axons. This was reflected in the quantification of the number of axons. Thickness of myelin sheaths, however, tended to be larger in the injured nerves (Supplementary Fig. 1), likely due to the expected separation of individual myelin layers and intra-myelin inclusions, as depicted by electron microscopic observation (Fig. 7 and Supplementary Fig. 4). Hypertrophic Schwann cell processes and sparse phagocytes bearing a foamy cytoplasm are commonly observed in the injured optic nerve (Fig. 2f). Thus, the decrease in the number of myelinated axons interspersed with hypertrophic glial processes and interstitial macrophage infiltration contributes to the overall atrophic appearance noted in the injured optic nerve. The number of axons was considerably lower as compared to intact nerves, reducing axonal density by around 50%, which translated into reductions of axon and myelin volume fractions (Fig. 4a,b and Supplementary Fig. 1). The remaining structurally viable axons in the injured nerves tended to have slightly larger calibers, with a proportionally larger loss of small-caliber axons (Fig. 7e). Sagittal sections of the intact optic chiasm showed two distinct, interdigitating axonal populations (Fig. 2d). Comparable sections from experimental animals showed one normally-appearing axonal population intermingled with collections of collapsed axons (Fig. 2h).

**Figure 2:**
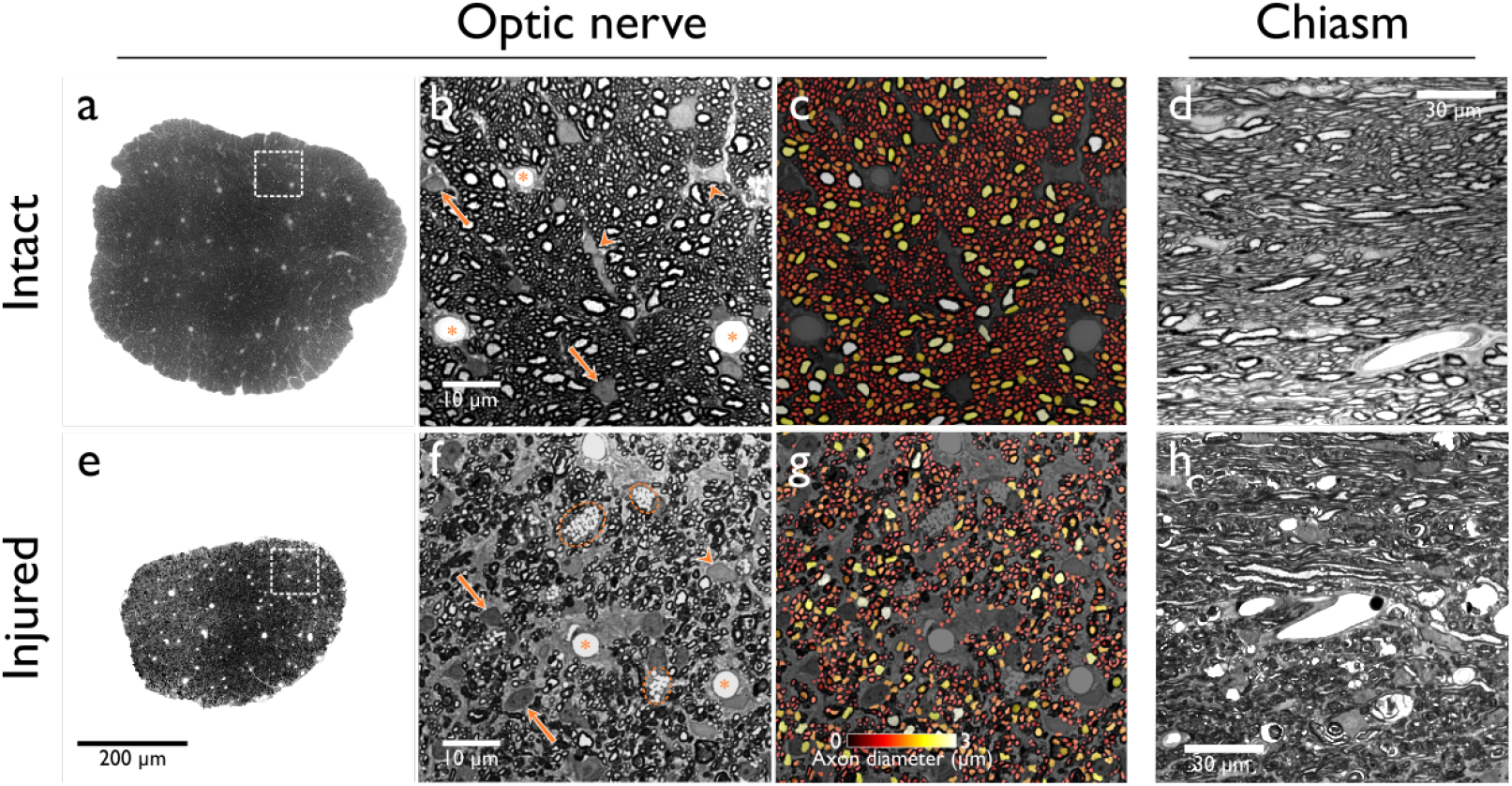
Histological evaluation of intact and injured optic nerves. As compared with the intact optic nerve (a), the injured nerve (e) shows considerable atrophy at low magnification. Higher magnification shows tightly-packed axons with dense, circular myelin sheaths in the intact nerve (b). Interstitial Schwann cells (arrows) and their processes (arrow heads), some of them encircling capillaries (asterisks), are observed. The damaged optic nerve (f) shows a dramatic reduction of normally-appearing axons, which are substituted by hyperchromatic axons with shrunken axoplasm and retracted myelin sheaths. Glial processes (arrows) are seemingly increased in size and number, and foamy phagocytes (dashed) are also frequent. Axon segmentation shows a marked decrease of axons in the injured nerve (g) compared to the intact optic nerve (c). The intact optic chiasm (d) shows two axonal populations running in different directions. In the injured optic chiasm (h) there are numerous clusters of altered axons that resemble those in panel (f). Notably, clusters of altered axons alternate with axons of normal appearance, most of them obliquely sectioned.

*Ex vivo* dMRI of the optic nerves and chiasm provided a high level of detail in both structures (Fig. 3). All three dMRI analysis methods were able to convey relevant information regarding the intact and injured nerves and chiasm (Fig. 4c-h and Supplementary Fig. 2). The CSD-derived FODs within the intact optic nerves showed a single fiber population and had large amplitudes. At the level of the intact chiasm, two fiber populations crossing nearly orthogonally can be seen. Both multi-tensor methods showed single-fiber populations with very high anisotropy in intact optic nerves, and two high-anisotropy tensors within the optic chiasms; those derived from the DIAMOND model were more anisotropic than those derived from MRDS. Injured optic nerves showed reduced amplitude of FODs and the presence of spurious peaks, as well as reduced anisotropy in multi-tensor models. Injured chiasms showed considerable amplitude reduction of the lobes of the FODs corresponding to the injured axonal population, whereas MRDS and DIAMOND showed reduced anisotropy or reduced compartment fraction of the tensors related to the injured bundle. Statistical pair-wise comparisons for the different metrics derived from the three methods are shown in Supplementary Fig. 2.

**Figure 3:**
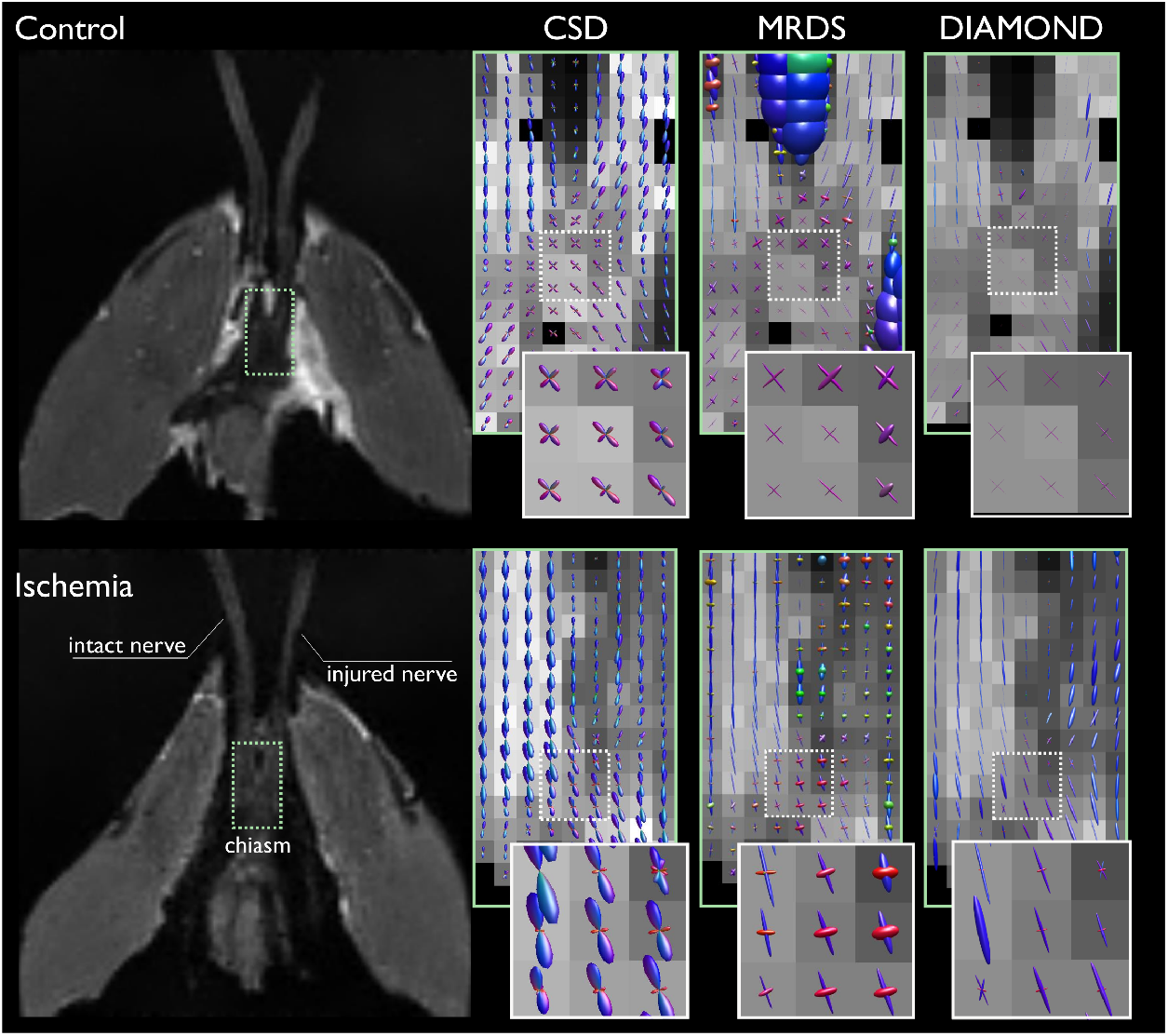
Qualitative evaluation. Analysis of a control specimen (top) shows both optic nerves having single-fiber FODs with large amplitude, as evaluated through CSD; these single-fiber FODs correspond to high anisotropy of single tensors for both multi-tensor models (MRDS and DIAMOND). The intact optic chiasm shows two fiber populations with nearly identical diffusion properties crossing the midline. Unilateral retinal ischemia (bottom) induced a low amplitude of FODs and low anisotropy in the injured nerve, whereas the optic chiasm shows a low amplitude of the FODs corresponding to the injured nerve (those from the intact nerve have a high amplitude), and low anisotropy (MRDS) and low volume fraction (DIAMOND). The dotted green and white regions are enlarged and framed in the same colors.

**Figure 4:**
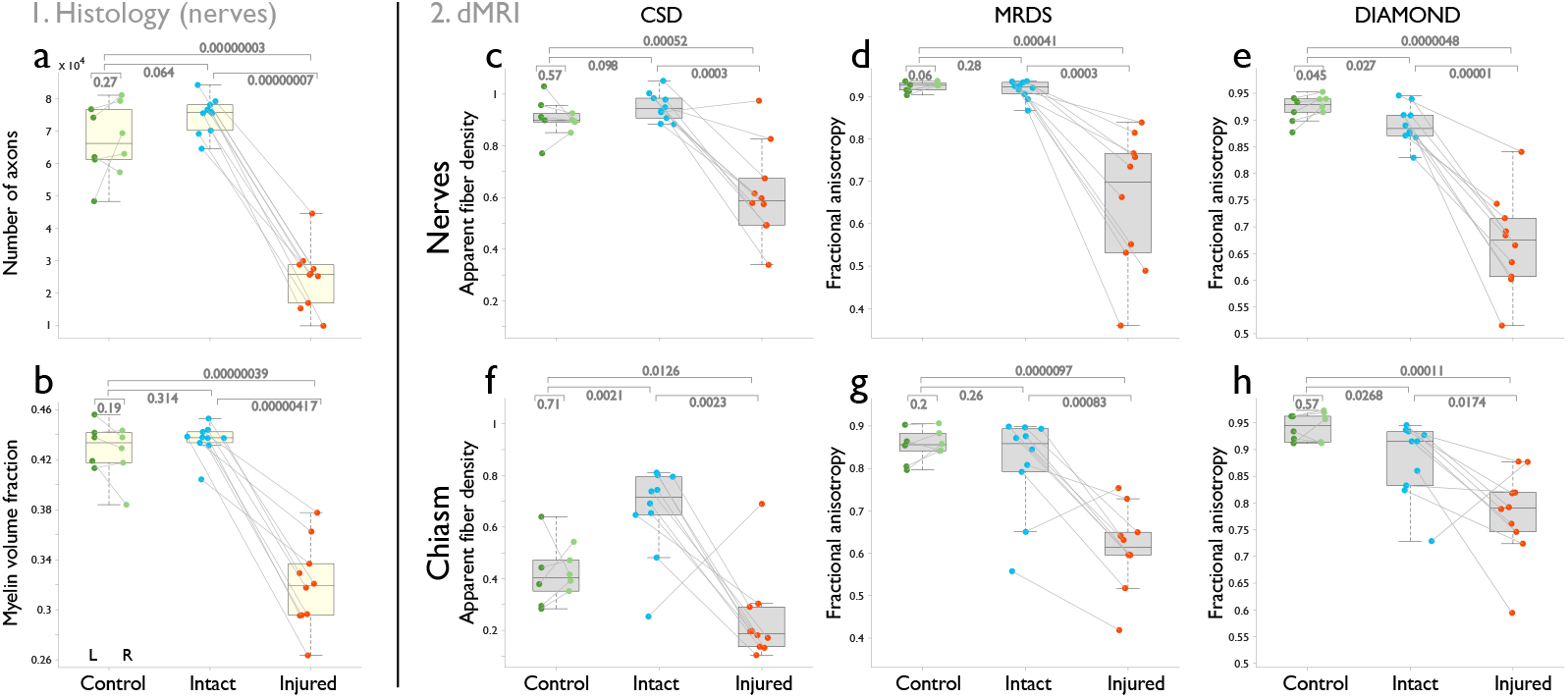
Main parameters derived from quantitative histology and dMRI of optic nerves and chiasm. The number of axons is drastically reduced in the injured optic nerves (a), which is accompanied by a reduction of myelin volume fraction (b). The three multi-fiber dMRI metrics clearly differentiate intact from injured nerves as reduced AFD (CSD), and FA (MRDS and DIAMOND) (panels c-e). In the chiasm, both multi-tensor models showed reduced FA for the affected fiber bundle (g, h). CSD showed a reduction of AFD for the injured bundle in the chiasm and, unexpectedly, increased AFD of the intact bundle (f). Detailed histology and dMRI metrics are provided in Supplementary Figs. 1 and 2.

Metrics derived from the same dMRI data sets using three different analytical methods showed tight correlations with quantitative histology of the optic nerves and chiasm (Fig. 5; for an interactive version, visit https://doi.org/10.5281/zenodo.2574201). For CSD, the highest positive correlations were observed for AFD and peak AFD and axon density, AVF, MVF and g-ratio, Notably, negative correlations were seen between these two metrics and axon diameter and myelin thickness. This pattern was similar for dMRI data derived from intact and injured optic nerves and chiasm, albeit correlations were slightly lower in the chiasm. FOD dispersion did not correlate with any of the histological parameters in the optic nerves, but did so in the chiasm (with a similar pattern to the one previously described). FOD complexity correlated negatively with axon density, AVF, MVF and g-ratio, and positively with axon diameter and myelin thickness in the nerves, but not in the chiasm, where it had moderate positive correlations with AVF, MVF, g-ratio and axon diameter. Of note, FOD complexity is very low in intact optic nerves, and high in the intact chiasm, and the opposite in damaged structures (visualized in Fig. 3). The tensors derived from MRDS showed clear correlations with histology in the optic nerves, namely positive for FA, MD, λ_⊥_, and compartment size (*α*), with axon density, AVF, MVF and g-ratio; and negative for axon diameter and myelin thickness. λ_⊥_ showed the opposite direction of correlations with the same histological parameters. Similar yet slightly lower correlations were seen for DIAMOND, which also captured significant variance of histology-derived metrics in the heterogeneity indices of λ_║_ and λ_⊥_ (λ_║*HEI*_ and *λ*_⊥*HEI*_, respectivey). Per-bundle tensors identified by MRDS in the chiasm showed correlations with the histological characteristics of the corresponding optic nerves, particularly for FA, λ_⊥_, λ_║_ and *α*, but MD did not show significant correlations with histology. Similar to MRDS, per-bundle FA, λ_║_ and *α* derived from DIAMOND correlated with histology; contrary to the correlations seen in the optic nerves, heterogeneity indices did not correlate with histology.

**Figure 5:**
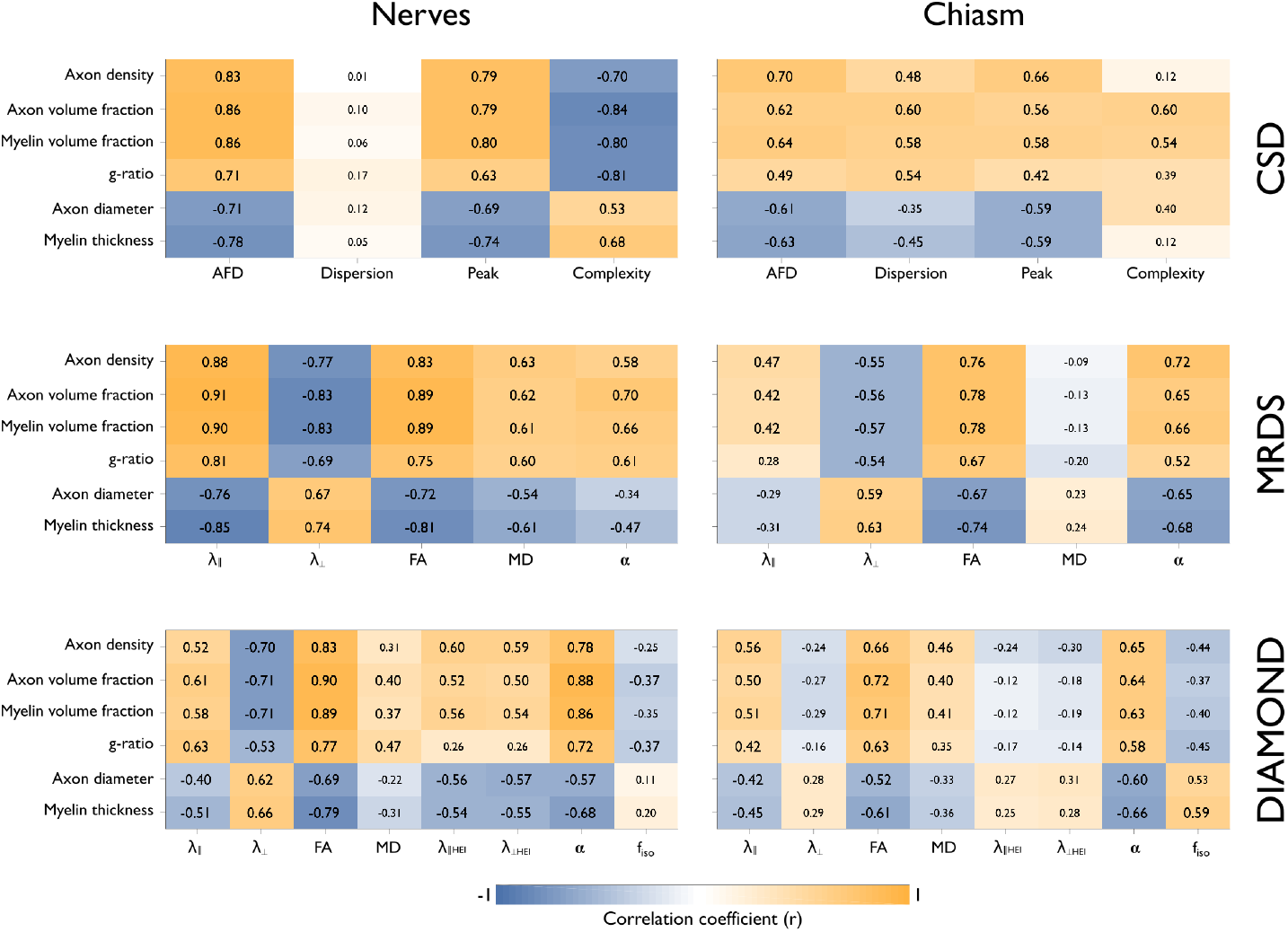
Correlation matrices between histological features (rows) and dMRI-derived metrics (columns) for the three dMRI methods used. In the optic neves, AFD, AFD peak, λ_║_, FA, MD and compartment size (*α*) show positive correlations with axon density, axon volume fraction, myelin volume fraction (expanded in Fig. 6), and g-ratio, and negative correlations with axon diameter and myelin thickness (see Fig. 7). This correlation pattern is similar for the corresponding bundles within the chiasm. The two multi-tensor models showed similar correlations with histology in the optic nerves for λ_║_, λ_⊥_, FA and *α*. Axial and radial heterogeneity (*H*_║_ and *H*_⊥_, respectively) of nerves, but not chiasm, correlated with histology. Contrary to MRDS, DIAMOND’s λ_║_ did not correlate with histology in the chiasm, whereas MD did. Each cell shows Pearson’s correlation coefficient, with bold typeface for *p* < 0.05. For a full list of p-values, see Supplementary Fig. 3.

Fig. 6 shows the positive correlations between axonal density and MVF with AFD and FA derived from dMRI for the optic nerves and chiasm. A continuum of histological alterations is observed for the control, intact and injured nerves that is closely mirrored by diffusion metrics. Segmentation of the FODs in CSD, and the estimation of two tensors by MRDS and DIAMOND at the level of the chiasm allows the estimation of dMRI metrics for each axonal population (i.e., fixel-wise analysis). This resulted in strong correlations that mimic those seen in the single-fiber population architecture of the optic nerves. Of note, while experimental animals showed the expected difference of AFD between the two bundles in the chiasm, the intact bundle showed values that were significantly higher than those seen in control animals (Figs. 4f and 6b, and Supplementary Fig. 2).

**Figure 6:**
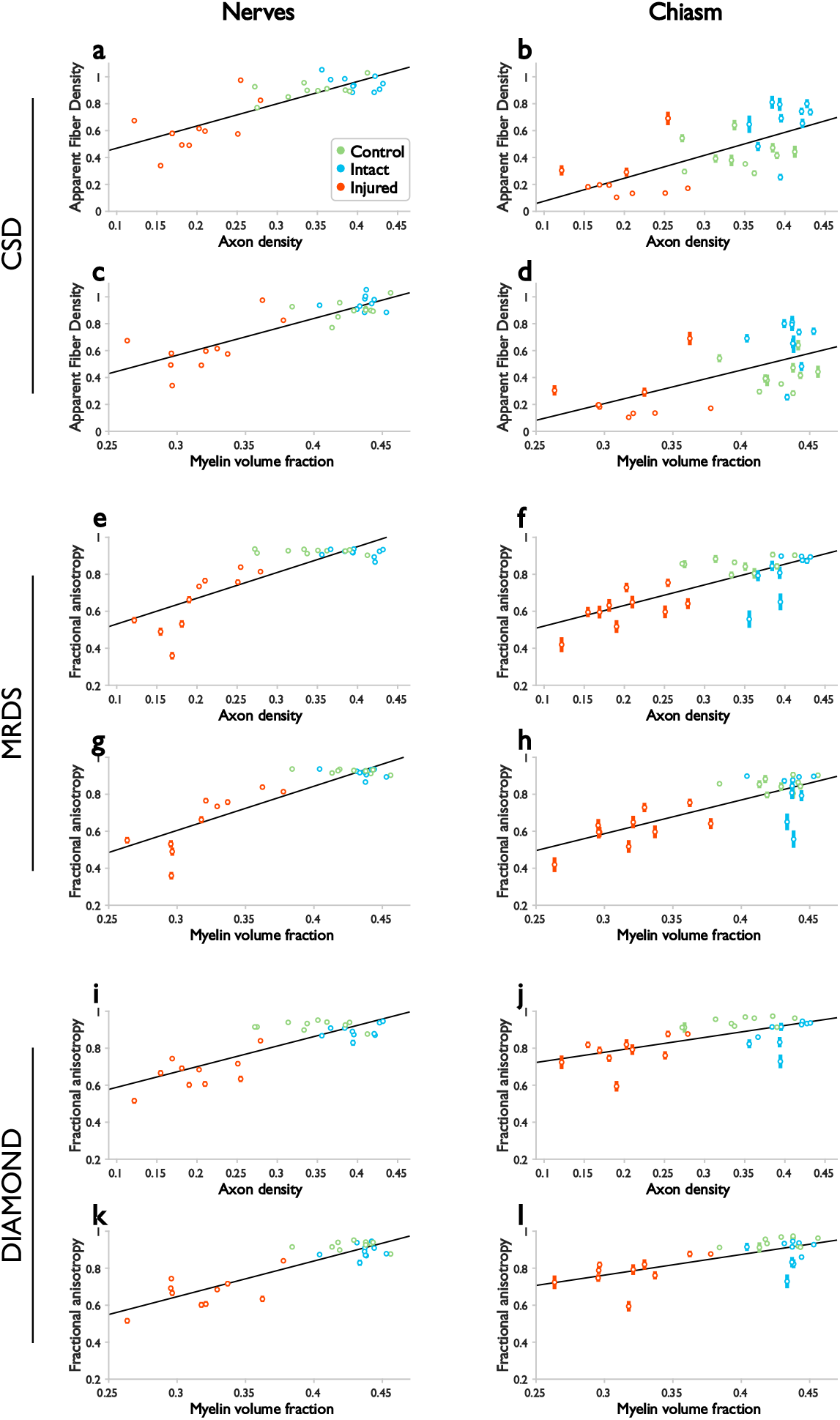
Correlations between dMRI metrics and quantitative histology. Strong positive correlations were observed for axon density and myelin volume fraction with AFD and FA in optic nerves and chiasm. The dMRI measurements are shown as mean (circles) and standard deviation (bars).

Negative correlations were found between axon diameter and myelin thickness, and diffusion metrics such as AFD and FA (Fig. 7). These negative correlations were unexpected and counterintuitive. Close examination of the automatic segmentation of axons and myelin sheaths provided insight into their genesis. First, while it is evident that axonal density is dramatically reduced in the injured nerves, as compared to their intact counterparts (Fig. 2), there seems to be a slight preferential loss of small-caliber axons, as can be seen in the joint histogram of all photomicrograph mosaics (Fig. 7e). Second, myelin thickness tends to be overestimated by the automatic segmentation method used here, particularly in the presence of intramyelin inclusions and separation of myelin layers. These phenomena are subtle and easily missed by the segmentation method when working on light microscopy images, but conspicuous and frequent when visualized through electron microscopy of the same specimens (Fig. 7k, and Supplementary Fig. 4).

**Figure 7:**
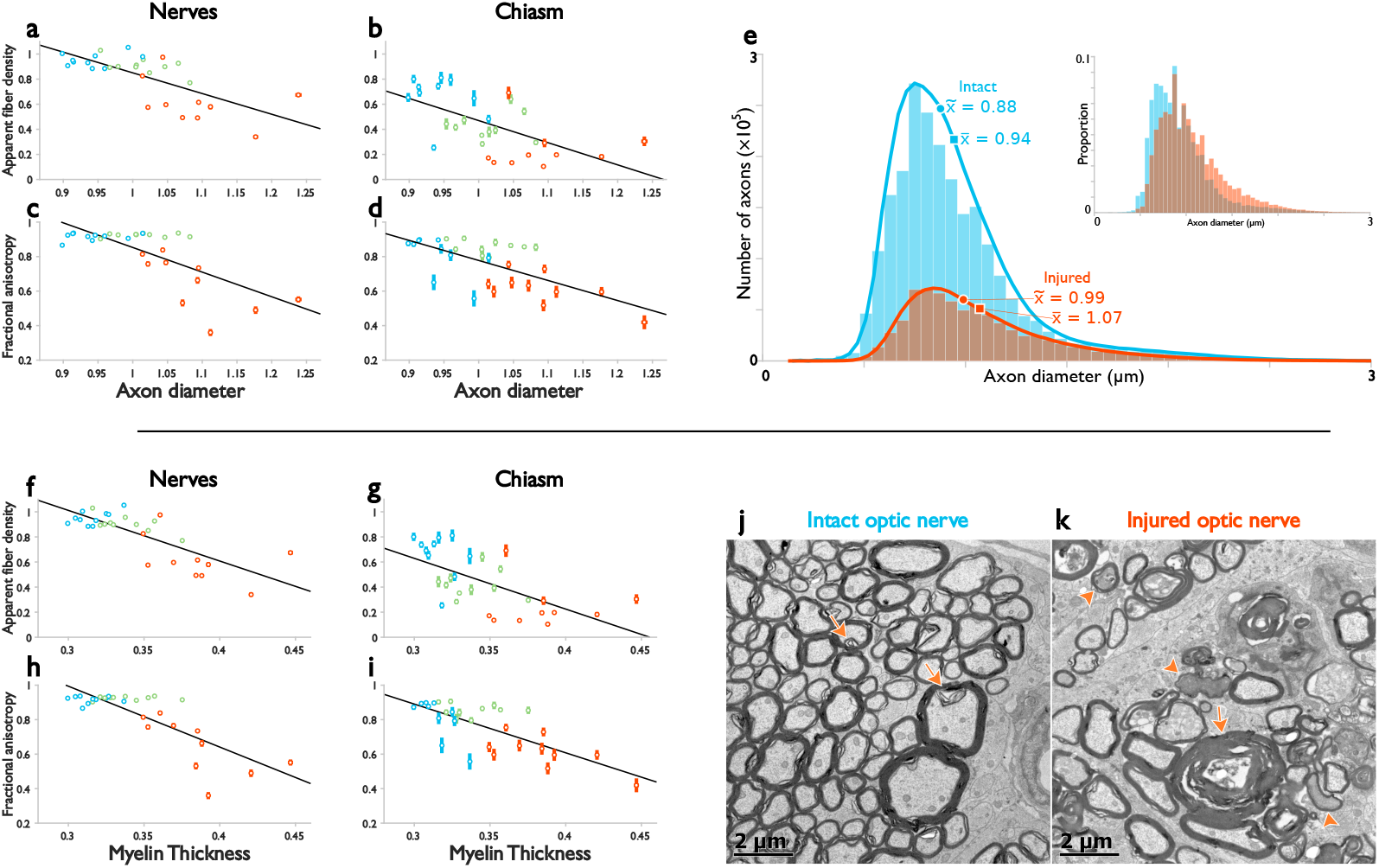
Correlations between dMRI metrics and quantitative histology. Negative correlations were observed for AFD and FA with average axon diameter (a-d). These correlations appear to be driven by preferential loss of small-caliber axons, as evidenced by the histogram resulting from the concatenation of all axons in all photomicrograph mosaics (e). Myelin thickness also correlated negatively with AFD and FA in both optic nerves and chiasm (f-i). This negative correlation is likely artifactual, resulting from the overestimation of myelin thickness when sheaths are separated (arrows in [j] and [k]), a phenomenon that is exaggerated in injured nerves (k). Arrowheads in (k) show degenerated axons with electro-dense cytoplasm. Panels (j) and (k) are exemplary survey electron micrographs of optic nerves at higher magnification than that used for automatic segmentation of axons. In the intact optic nerve (j) most axons bare compact myelin envelopes. Myelin splitting is occasionally observed in the normal optic nerve (arrows), but it occurs more extensively and frequently in the injured optic nerve (k), which yields an increased overall myelin thickness. FA shown in panels (c), (d), (h), and (i), is derived from MRDS; similar correlations are observed for DIAMOND (not shown).

The results of the BAM analysis, shown in Fig. 8, display the robustness of the correlations between dMRI descriptors and metrics computed from histology when the number of DWI acquisitions is reduced. These results corroborate that many of the dMRI descriptors are sufficiently robust to describe histological features, even when the number of volumes is reduced considerably (from 80 to 30 DWI volumes per shell). The most reliable estimators (i.e., those with consistent correlation coefficients obtained from different numbers of DWI volumes) for nerves and chiasm included CSD’s AFD and peak AFD, MRDS’s FA, λ_⊥_ and *α*, and DIAMOND’s FA and *α*.

**Figure 8:**
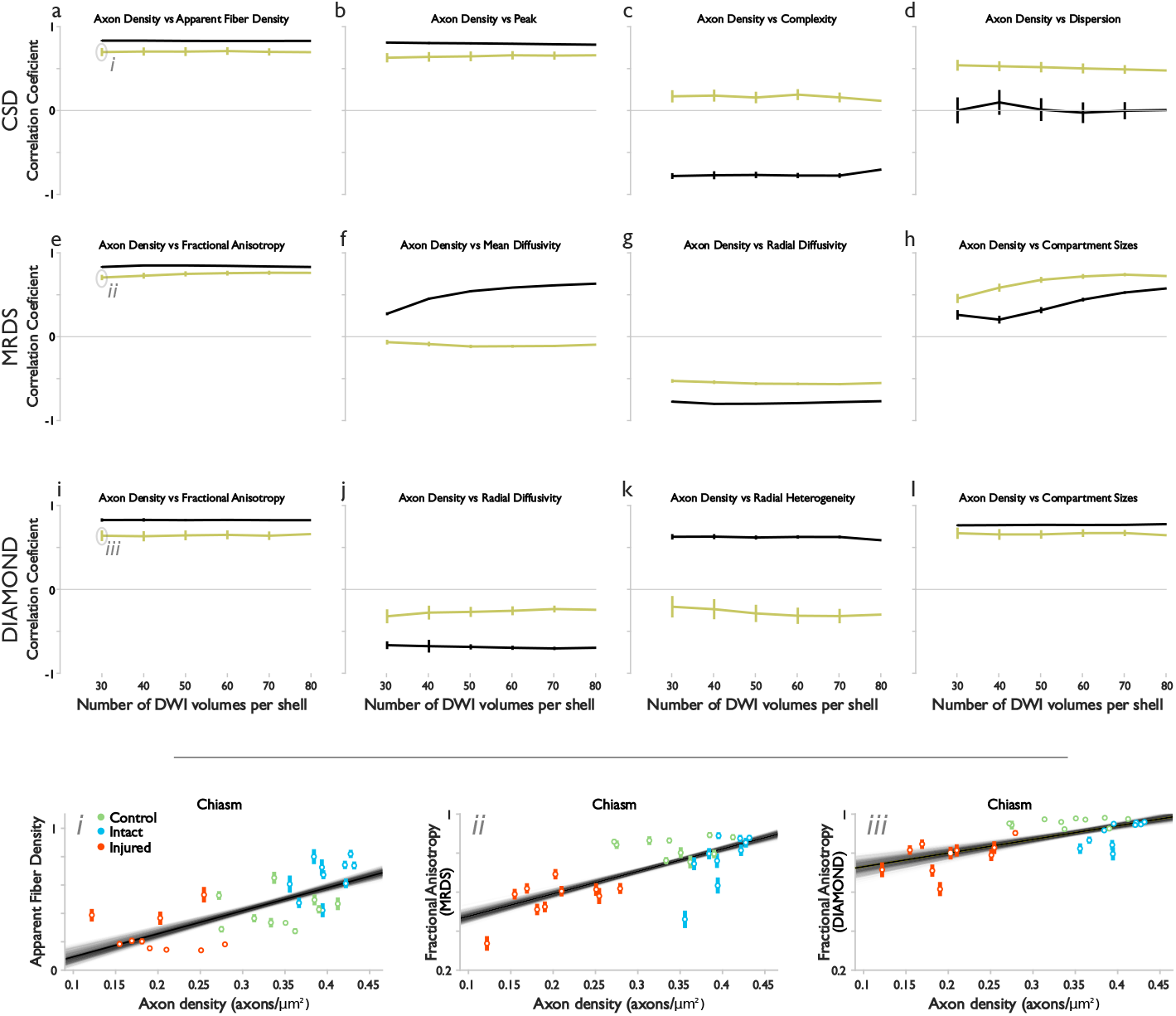
Robustness of correlations with respect to the number of volumes in the data set. Out of the 160 DWI volumes available in the original data set (80 DWI directions in two shells), we evaluated correlations with histology using a subset of *n* = [30 : 10 : 70] randomly selected volumes per shell. Shown here are only correlations with axon density. The top panel shows stability of correlation coefficients as a function of number of DWI volumes as mean values (horizontal lines) and standard deviations (vertical lines). Most correlations are very stable regardless of the number of volumes used (n ≥ 30/shell). Exemplary bagging results identified as *i*, *ii* and *iii* in (a), (e), and (i), respectively, are expanded in the bottom panel, where the overlapping semi-transparent lines indicate the resulting linear regression for a particular bagging realization.

## 4. Discussion

This work provides direct evidence of the ability of three different dMRI processing methods to capture relevant microstructural information related to white matter damage in regions fiber crossing regions. We extended the well-known model of unilateral retinal ischemia (Adachi et al., 1996; Song et al., 2003) to induce Wallerian degeneration of the optic nerve emerging from the affected eye, providing an ideal opportunity to evaluate the optic chiasm, where the degenerated axons cross their intact counterparts emanating from the contralateral (unaffected) eye. The known anatomy of the visual pathway and the discrete nature of the induced white matter damage allowed us to evaluate the extent to which quantitative parameters derived from advanced dMRI models relate to tissue characteristics.

Axonal degeneration produces morphological changes that are readily followed by dMRI in regions where a single fiber population exists (Beaulieu et al., 1996; Stanisz et al., 2001). The tensor model is able to distinguish the acute and chronic stages of Wallerian degeneration, characterized by an initial fragmentation of axons, followed by myelin degradation and glial infiltration (Song et al., 2003; Concha et al., 2006; Liu et al., 2013). The non-invasive identification and staging of this pathological phenomenon is invaluable in clinical and research settings. However, such a feat is not trivial in regions with complicated fiber architecture, such as in voxels containing crossing, fanning, kissing or incoherently-organized fibers (Tournier et al., 2011). With most of the human white matter containing such complex configurations (Jeurissen et al., 2013), the need to disentangle fiber populations and derive their characteristics independently is obvious. Yet, in addition to their sensitivity, metrics derived from dMRI should be easily relatable to biological features of the kind a neuro-pathologist may describe.

In the optic nerves, we found the expected morphological changes that characterize the chronic phase of Wallerian degeneration, with the most striking features being a profound loss of axonal density, severe degradation of myelin, and gross atrophy (Fig. 2). The single tensor model also showed the expected reduction of diffusion anisotropy at the expense of increased λ_⊥_. AFD was significantly reduced in degenerated optic nerves, and the emergence of spurious peaks in the resulting FODs abnormally increased their complexity (Fig. 3). The three dMRI analysis methods correctly identified the two fiber populations contained in the intact optic chiasm. Notably, degeneration of one of the fiber populations in this crossing region was evidenced by a reduction of the amplitude of the corresponding lobe in the FOD (i.e., reduced AFD). The tensor identified as belonging to the degenerated fibers showed reduced FA and volume fraction, and increased heterogeneity and λ_⊥_, analogous to the injured optic nerves.

Strong correlations were found between dMRI metrics and histological features. The most relevant is the positive association between axonal density and FA and AFD. Importantly, these correlations are evident at the level of the optic nerves and chiasm (Fig. 5). Thus, when multi-fiber reconstructions are performed, these two metrics can be interpreted in a similar fashion in regions of one or more axonal populations. However, other metrics should be analyzed with caution. FOD complexity, for example, is very low in intact optic nerves and high in the intact optic chiasm, but in the presence of degenerated axons, this pattern is reversed. This finding is in line with expectations and the appearance of resulting FODs (Fig. 3). However, we unexpectedly found that myelin thickness and axon diameter correlated negatively with the AFD and FA of the nerves and chiasm (Fig. 7). Close examination using electron microscopy revealed that myelin thickness was overestimated by the method used for automatic segmentation because of the abundance of intra-myelin inclusions and fraying of myelin sheaths (Fig. 7-j,k and Supplementary Fig. 4). Some intramyelin inclusions and myelin sheath separations are even seen in intact optic nerves (Fig. 2), which are expected given the relatively long time between tissue fixation and preparation for histology (Peters, 1970; Hukkanen and Röyttä, 1987). However, the degree of myelin abnormalities is considerably greater in the injured nerves. The artifactual negative correlation between myelin thickness and AFD/FA is corrected when accounting for axonal density, in the form of myelin volume fraction, which correlated positively with AFD, FA, and MD (Fig. 5). Degenerated nerves showed that the small fraction of surviving axons had, on average, a slightly larger diameter than those of intact nerves. Inspection of the histogram of axonal diameters across all the samples revealed that large caliber axons were preferentially spared after injury (Fig. 7e). The different vulnerability of axons as a function of their diameter has been reported in post-mortem studies of spinal cords of patients with multiple sclerosis, although the mechanisms that drive this phenomenon are unclear (DeLuca et al., 2004). The contribution of very small caliber axons (< 0.2 *μ*m) to the correlations presented herein is also difficult to predict, since those axons cannot be resolved with optical imaging (Innocenti et al., 2013). Glial infiltration was observed in the degenerated nerves and chiasm. It has been reported that glial scars formed after spinal cord injury can create anisotropic structures perpendicular to axons, thereby modulating diffusion anisotropy in degenerated white matter bundles (Schwartz et al., 2005). We cannot comment on the influence of glial cells on dMRI and measured anisotropy, as we did not characterize the spatial organization of these cells. In the chiasm, AFD of the intact fiber bundle of experimental animals was larger than that of control animals (Fig. 6b and Supplementary Fig. 2), likely due to the use of a fixed response function in the CSD methodology. The signal from the injured bundle could not be fully represented by its corresponding FOD lobe, causing an increase in the lobe corresponding to the intact lobe in order to minimize the error in the model. This was corroborated using synthetic data and presented in Supplementary Fig. 5. Despite the ability of CSD to disentangle the intact and injured axonal populations, careful interpretation of absolute values of AFD is thus warranted in future studies using such metric.

The data sets in this study used 80 directions for diffusion sensitization each in two *b* > 0 s/mm^2^ shells. Even with rapid acquisition techniques, such as echo-planar imaging, the acquisition time would be too long for most clinical settings. We therefore tested if the correlations between dMRI metrics and histological features are viable with small data sets. Our results (Fig. 8) indicate that dMRI descriptors provide useful correlations about the healthy and damaged microstructure features, even with a reduced set of acquisitions. The majority of correlations we identified with the complete data set are also observed when using only 30 acquisitions per shell, which is attainable for clinical purposes. Our results indicate that the proposed animal model is useful to characterize the performance of methods for dMRI analysis. This biological model provides an excellent opportunity to evaluate the performance of methodologies that yield additional descriptors of tissue, such as intracellular volume fractions and axon diameter estimations (Assaf et al., 2008; Alexander et al., 2010; Fieremans et al., 2011; Zhang et al., 2012; Panagiotaki et al., 2012; Dyrby et al., 2013), and alternative image acquisition approaches (Mitra, 1995; Shemesh et al., 2016; Yang et al., 2018).

*Ex vivo* imaging provides the opportunity to acquire very high resolution images through long acquisition times, with substantial minimization of artifacts such as motion and magnetic susceptibility. However, there are crucial differences between *ex vivo* and *in vivo* tissue that can potentially confound the interpretation of dMRI. First, blood vessels no longer present flow; instead, they are occupied by the fixative solution used, and the fast and directional motion of blood is replaced by a large pool of diffusing water molecules. Given the relatively large caliber of vessels (as compared to axons, and in the context of the effective diffusion times used in this study), this water pool likely shows fast and isotropic diffusion. Although multi-tissue CSD attempts to capture a freely-diffusing compartment (Jeurissen et al., 2014), and DIAMOND estimates the isotropic fraction *f_iso_*, MRDS does not not explicitly search for this pool. Second, temperature directly affects dMRI. Temperature variations modulate diffusivity, which would confound anisotropy measures, as each diffusion-weighted volume would be differentially affected. In this study, room temperature was kept at 21¤C, thus minimizing this confound. Nonetheless, tissue temperature is lower than *in vivo*, thus reducing overall diffusivity, but not anisotropy (D’Arceuil et al., 2007). It is therefore recommended that the selected *b* values for *ex vivo* imaging be three to four times higher than those typically selected for *in vivo* applications (Dyrby et al., 2011). The *b* values used here (2000 and 2500 s/mm^2^) can thus be considered low. Despite this level of diffusion sensitization, we showed that three different dMRI analysis methods were able to precisely identify the crossing fibers in the chiasm, and accurately identify diffusion alterations related to axonal degeneration. This is encouraging, since most clinically-available scanners are equipped with low-amplitude gradient systems that would require long echo times to achieve high *b* values, negatively impacting signal-to-noise ratio (Jones et al., 2018). Lastly, tissue shrinks after perfusion fixation, and even more so after preparation for microscopy, mainly due to a reduction of the extra-cellular space (Dyrby et al., 2018). Thus, contributions to the dMRI signal from intra- and extra-axonal compartments may differ from what is appreciated through light and electron microscopy.

Experimental logistics of our study resulted in brains being imaged at different times with respect to euthanasia, and specimens processed in different batches. These factors potentially introduced biases in the estimation of dMRI metrics and quantitative histology (e.g., differences in axonal density of optic nerves between control animals and intact nerves in the experimental group; Supplementary Fig. 1). Another limitation is the lack of direct estimation of histological features in the chiasm, which were estimated based on the average of features obtained in the optic nerves. Given the intricate arrangement of axons within the chiasm (Colello and Guillery, 1998), automatic segmentation of axons was either inadequate or biased towards a particular population. Our approach to infer chiasm tissue characteristics based on those measured in the optic nerves relies on the fact that most retinal axons decussate in rodents (Jeffery and Erskine, 2005). Other forms of morphological analysis of this structure, such as those derived from large field-of-view, three-dimensional electron microscopy (Denk and Horstmann, 2004; Abdollahzadeh et al., 2018; Pichat et al., 2018; Lee et al., 2019) or optical coherence tomography (Lefebvre et al., 2018), should provide more accurate estimations of tissue properties, including many that cannot be resolved from two-dimensional sections, such as axon tortuosity and dispersion, and spatial arrangement of glial cells, vessels, and other structures that contribute to modulation of water diffusion.

Several studies have used histological methods to validate fiber orientations derived from dMRI (Wang et al., 2014; Sierra et al., 2015; Khan et al., 2015; Salo et al., 2018; Chang et al., 2017; Lefebvre et al., 2018), showing agreement between the two, even in regions of fiber crossings (Leergaard et al., 2010; Budde and Annese, 2013; Schilling et al., 2016; Axer et al., 2016). Yet, to be useful for clinical and research purposes, per-bundle diffusion characteristics are needed, along with their respective validation with respect to a gold standard, as has been largely achieved in the case of single-fiber regions (Concha, 2014; Alexander et al., 2017). Our work provides grounds for proper interpretation of diffusion abnormalities in fiber crossing regions. The three dMRI methods validated here were chosen because they have relatively lenient acquisition requirements, and provide metrics that are intuitively related to tissue characteristics. Our results show that these dMRI methods provide reliable indicators of white matter damage in regions with one or more axonal populations, even at relatively low *b* values and few dMRI volumes.

This work approaches diffusion metrics as biomarkers of broad characteristics of tissue (as opposed to their use in biophysical models that extract specific tissue parameters) (Jespersen, 2018), and investigates how much biologically-relevant information a specific dMRI representation captures (Novikov et al., 2018). Such an approach favors sensitivity over specificity, but may provide more tangible metrics that clinicians and researchers can use to diagnose and monitor various neurodegenerative disorders. At the same time, histological validations of dMRI metrics will minimize their overinterpretation and misinterpretation, which have unfortunately become all too common (Jones et al., 2013; Novikov et al., 2018). Accurate interpretation of dMRI, used in combination with other methods, will further improve the yield of detailed information on tissue microstructure in a non-invasive fashion (Cercignani and Bouyagoub, 2018). As new forms of analysis of the dMRI signal are developed, they should undergo rigorous validation through numerical simulations, physical phantoms and, ultimately, histological correspondence. To this end, we provide all the dMRI and histological data through the White Matter Microscopy Database (Cohen-Adad et al., 2017).

## Acknowledgements

Imaging was performed at the National Laboratory for Magnetic Resonance Imaging (Conacyt, UNAM, CIMAT, UAQ). We thank Juan Ortiz-Retana for technical assistance related to MRI; and Jessica González-Norris for proof-reading and editing our manuscript. We also thank Erick Pasaye-Alcaraz, Leopoldo González-Santos, Gema Martínez-Cabrera, and the personnel at the Animal Facility and Microscopy Unit of the Institute of Neurobiology, as well as the National Laboratory for Advanced Scientific Visualization, for technical support.

## Supplementary Material

**Supplementary Fig. 1:**
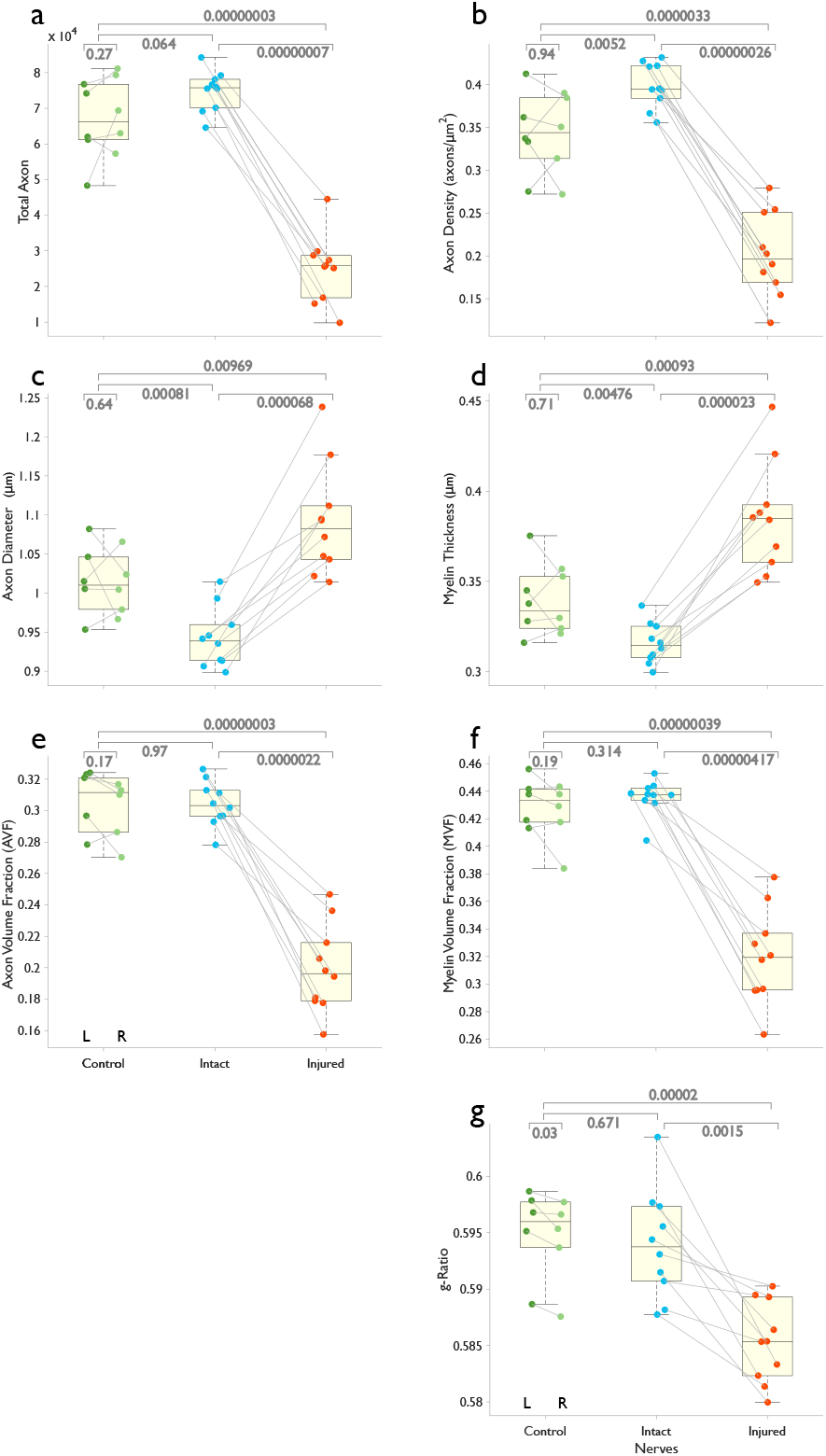
Quantitative histology of optic nerves. Each point represents the median value of each histological feature in three photomicrographs. Absolute axonal count and axonal density were greatly reduced in injured nerves following retinal ischemia. This reduction in the number of axons is reflected in considerable reductions of axonal and myelin volume fractions. Average axon diameter and myelin thickness increased slightly in injured nerves (thus reducing g-ratio), due to a preferential loss of small-caliber axons and an overestimation of myelin thickness caused by myelin sheath separation in injured nerves (see Figs. 2, 7, and Supplementary Fig. 4).

**Supplementary Fig. 2:**
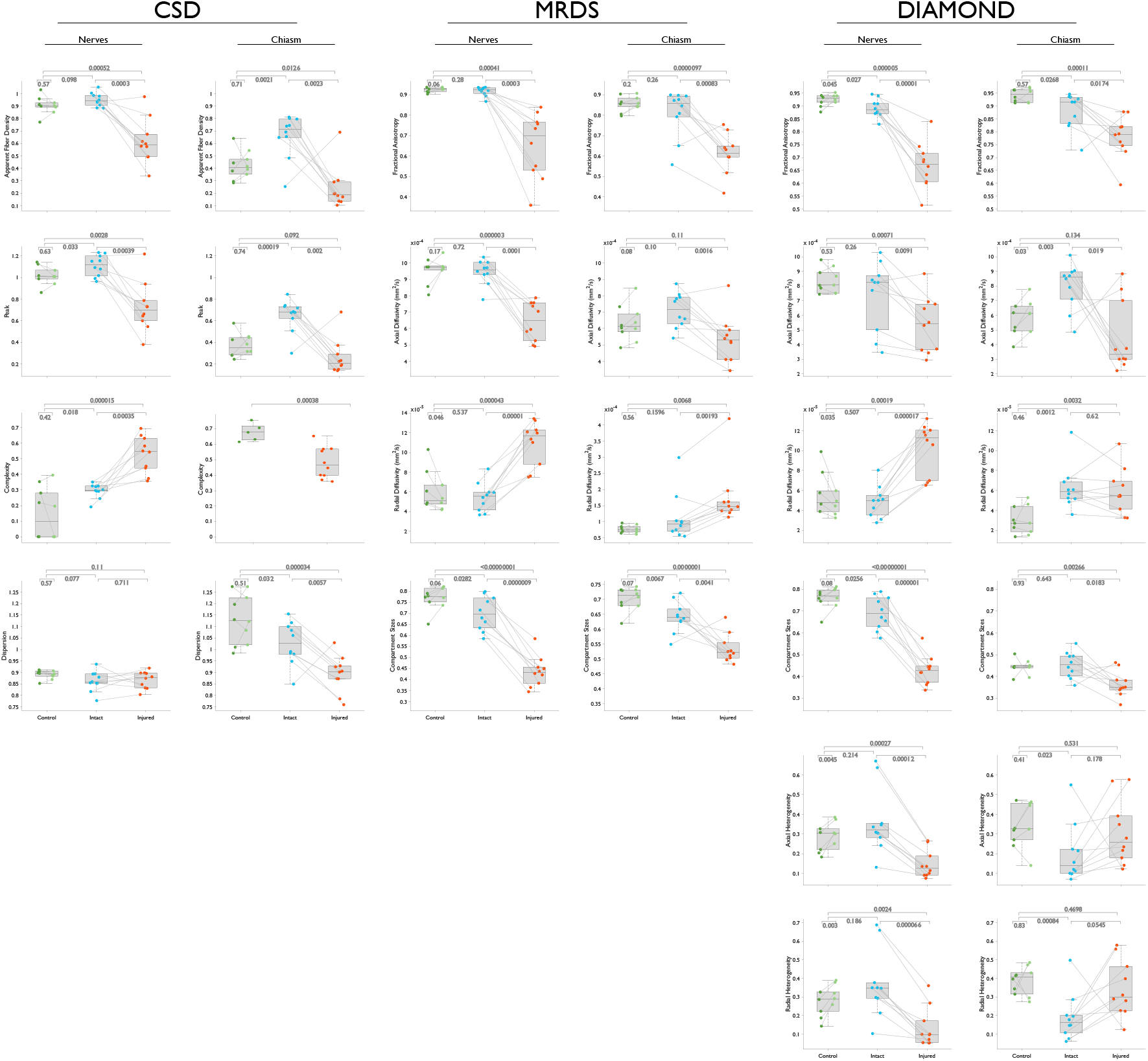
Pair-wise statistical tests for metrics derived from three dMRI methods. Paired t-tests for comparisons between left/right and injured/intact optic nerves or bundles in the chiasm. Independent samples t-tests for between-group comparisons.

**Supplementary Fig. 3:**
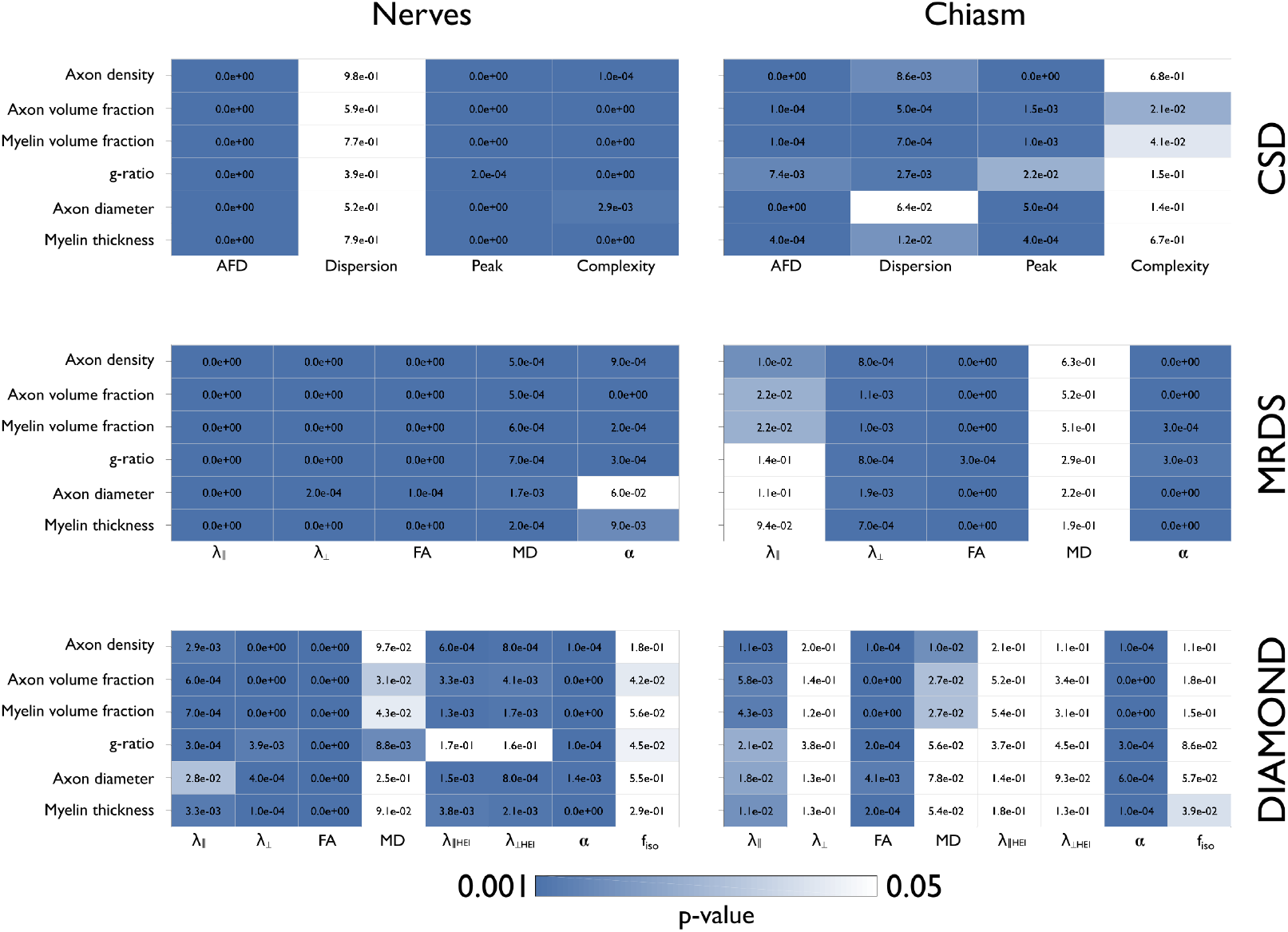
P-values corresponding to the correlation matrices between histology-based values and dMRI-derived metrics shown in Fig. 5. 10,000 permutations.

**Supplementary Fig. 4:**
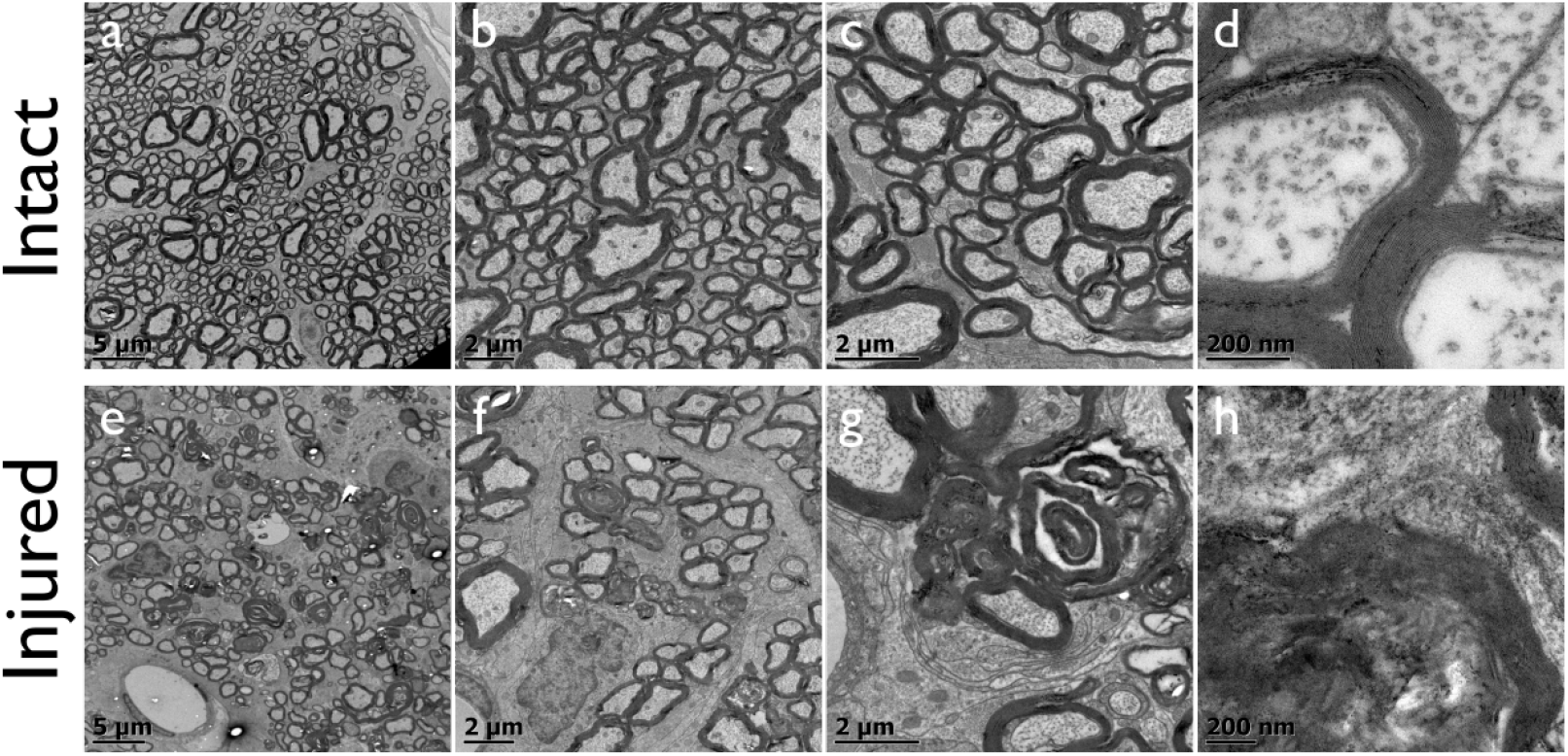
Electron micrographs of intact and injured optic nerves at increasingly higher magnifications.

**Supplementary Fig. 5:**
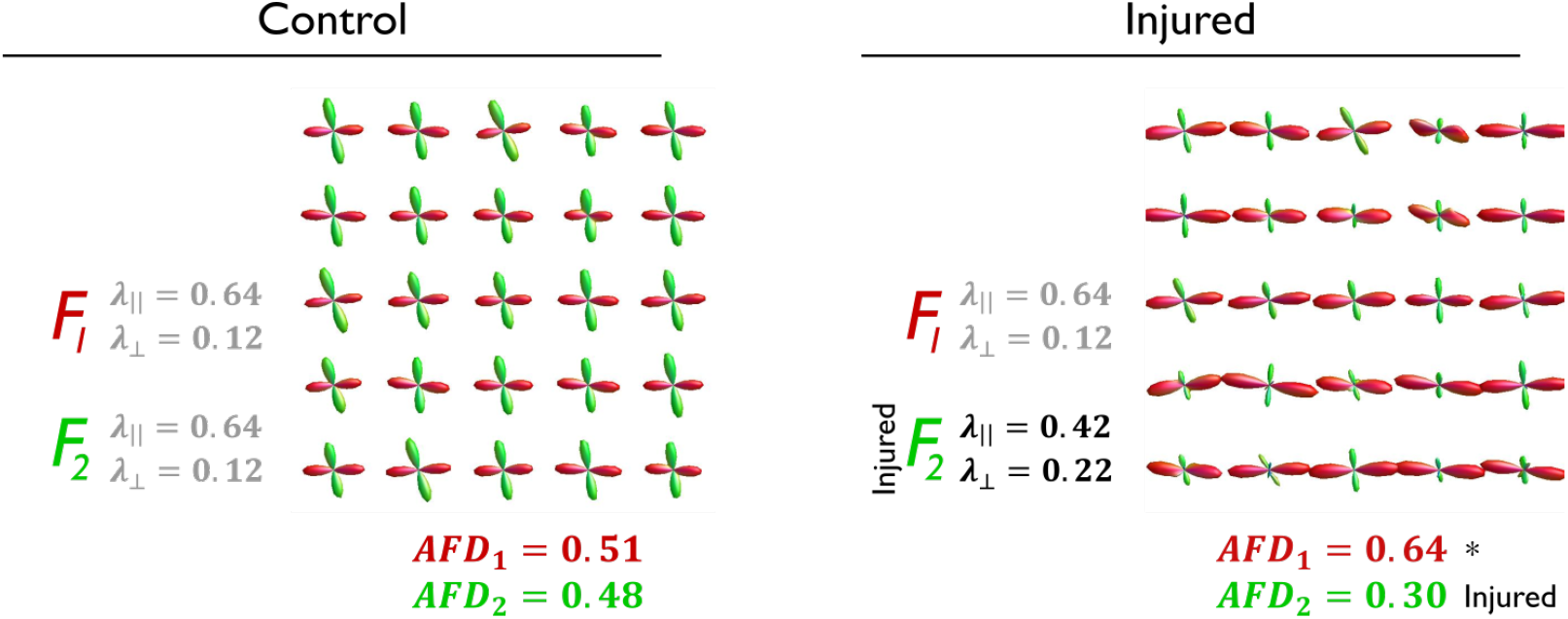
Artifactually increased AFD in fiber crossing regions in the presence of degeneration. Synthetic data show two fiber populations (*F*_1_,*F*_2_), each with *α* = 0.5, crossing at nearly right angles. *F*_1_ and *F*_2_ have similar λ_║_ and λ_⊥_ in the control condition, while injury to *F*_2_ is simulated by reducing its λ_║_ and increasing λ_⊥_. The resulting AFD (averaged over all FODs) is nearly identical for both fiber bundles in the control condition. In the injured case, AFD for *F*_2_ is reduced, albeit with an unexpected increase of AFD for *F*_1_ (asterisk). λ_║_ and λ_⊥_ with units ×10^−3^ mm^2^/s.

## References

Abdollahzadeh, A., Belevich, I., Jokitalo, E., Tohka, J., Sierra, A., 2018. 3d axonal morphometry of white matter. bioRxiv, 239228.

Adachi, M., Takahashi, K., Nishikawa, M., Miki, H., Uyama, M., 1996. High intraocular pressure-induced ischemia and reperfusion injury in the optic nerve and retina in rats. Graefe’s Archive for Clinical and Experimental Ophthalmology = Albrecht Von Graefes Archiv Für Klinische Und Experimentelle Ophthalmologie 234, 445–451.

Akaike, H., 1974. A new look at the statistical model identification, in: Selected Papers of Hirotugu Akaike. Springer, pp. 215–222.

Alexander, D.C., 2005. Multiple-fiber reconstruction algorithms for diffusion mri. Annals of the New York Academy of Sciences 1064, 113–133.

Alexander, D.C., Dyrby, T.B., Nilsson, M., Zhang, H., 2017. Imaging brain microstructure with diffusion mri: practicality and applications. NMR in Biomedicine, e3841.

Alexander, D.C., Hubbard, P.L., Hall, M.G., Moore, E.A., Ptito, M., Parker, G.J., Dyrby, T.B., 2010. Orientationally invariant indices of axon diameter and density from diffusion {MRI}. NeuroImage 52, 1374–1389.

Assaf, Y., Blumenfeld-Katzir, T., Yovel, Y., Basser, P.J., 2008. Axcaliber: A method for measuring axon diameter distribution from diffusion mri. Magnetic Resonance in Medicine 59, 1347–1354.

Axer, M., Strohmer, S., Gräßel, D., Bücker, O., Dohmen, M., Reckfort, J., Zilles, K., Amunts, K., 2016. Estimating fiber orientation distribution functions in 3d-polarized light imaging. Frontiers in neuroanatomy 10, 40.

Basser, P.J., Mattiello, J., LeBihan, D., 1994. MR diffusion tensor spectroscopy and imaging. Biophysics Journal 66, 259–67.

Beaulieu, C., 2002. The basis of anisotropic water diffusion in the nervous system - a technical review. NMR in Biomedicine 15, 435–55.

Beaulieu, C., Does, M.D., Snyder, R.E., Allen, P.S., 1996. Changes in water diffusion due to wallerian degeneration in peripheral nerve. Magnetic resonance in medicine 36, 627–631.

Breiman, L., 1996. Bagging predictors. Machine Learning 24, 123–140.

Budde, M.D., Annese, J., 2013. Quantification of anisotropy and fiber orientation in human brain histological sections. Frontiers in Integrative Neuroscience 7, 3.

Budde, M.D., Kim, J.H., Liang, H.F., Schmidt, R.E., Russell, J.H., Cross, A.H., Song, S.K., 2007. Toward accurate diagnosis of white matter pathology using diffusion tensor imaging. Magn Reson Med 57, 688–695.

Cercignani, M., Bouyagoub, S., 2018. Brain microstructure by multi-modal mri: Is the whole greater than the sum of its parts? Neuroimage 182, 117–127.

Cercignani, M., Dowell, N.G., Tofts, P.S., 2018. Quantitative MRI of the brain: principles of physical measurement. CRC Press.

Chang, E.H., Argyelan, M., Aggarwal, M., Chandon, T.S.S., Karlsgodt, K.H., Mori, S., Malhotra, A. K., 2017. The role of myelination in measures of white matter integrity: combination of diffusion tensor imaging and two-photon microscopy of clarity intact brains. Neuroimage 147, 253–261.

Cohen-Adad, J., Does, M., Duval, T., Dyrby, T., Fieremans, E., Foias, A., Zaimi, A., 2017. White matter microscopy database. Open Sci. Framew. .

Colello, S., Guillery, R., 1998. European neuroscience association the changing pattern of fibre bundles that pass through the optic chiasm of mice. European Journal of Neuroscience 10, 3653–3663.

Concha, L., 2014. A macroscopic view of microstructure: Using diffusion-weighted images to infer damage, repair, and plasticity of white matter. Neuroscience 276, 14–28. Bibtex: concha_macroscopic_2014.

Concha, L., Gross, D.W., Wheatley, B.M., Beaulieu, C., 2006. Diffusion tensor imaging of time-dependent axonal and myelin degradation after corpus callosotomy in epilepsy patients. Neuroimage 32, 1090–1099.

Concha, L., Livy, D.J., Beaulieu, C., Wheatley, B.M., Gross, D.W., 2010. In vivo diffusion tensor imaging and histopathology of the fimbria-fornix in temporal lobe epilepsy. Journal of Neuroscience 30, 996–1002.

Coronado-Leija, R., Ramirez-Manzanares, A., Marroquin, J.L., 2017. Estimation of individual axon bundle properties by a Multi-Resolution Discrete-Search method. Medical Image Analysis 42, 26–43.

Daducci, A., Canales-Rodriguez, E., Descoteaux, M., Garyfallidis, E., Gur, Y., Lin, Y.C., Mani, M., Merlet, S., Paquette, M., Ramirez-Manzanares, A., Reisert, M., Reis Rodrigues, P., Sepehrband, F., Caruyer, E., Choupan, J., Deriche, R., Jacob, M., Menegaz, G., Prckovska, V., Rivera, M., Wiaux, Y., Thiran, J.P., 2014. Quantitative Comparison of Reconstruction Methods for Intra-Voxel Fiber Recovery From Diffusion MRI. IEEE Transactions on Medical Imaging 33, 384–399.

Daducci, A., Canales-Rodríguez, E., Zhang, H., Dyrby, T., Alexander, D., Thiran, J.P., 2015. Accelerated microstructure imaging via convex optimization (amico) from diffusion mri data. NeuroImage 105C, 32–44.

Dell’Acqua, F., Simmons, A., Williams, S.C., Catani, M., 2013. Can spherical deconvolution provide more information than fiber orientations? hindrance modulated orientational anisotropy, a true-tract specific index to characterize white matter diffusion. Human brain mapping 34, 2464–2483.

Dell’Acqua, F., Catani, M., 2012. Structural human brain networks: hot topics in diffusion tractography. Current opinion in neurology 25, 375–383.

DeLuca, G., Ebers, G., Esiri, M., 2004. Axonal loss in multiple sclerosis: a pathological survey of the corticospinal and sensory tracts. Brain 127, 1009–1018.

Denk, W., Horstmann, H., 2004. Serial block-face scanning electron microscopy to reconstruct three-dimensional tissue nanostructure. PLoS biology 2, e329.

Douaud, G., Jbabdi, S., Behrens, T.E., Menke, R.A., Gass, A., Monsch, A.U., Rao, A., Whitcher, B., Kindlmann, G., Matthews, P.M., et al., 2011. Dti measures in crossing-fibre areas: increased diffusion anisotropy reveals early white matter alteration in mci and mild alzheimer’s disease. Neuroimage 55, 880–890.

Dyrby, T.B., Baare, W.F., Alexander, D.C., Jelsing, J., Garde, E., Søgaard, L.V., 2011. An ex vivo imaging pipeline for producing high-quality and high-resolution diffusion-weighted imaging datasets. Human brain mapping 32, 544–563.

Dyrby, T.B., Innocenti, G.M., Bech, M., Lundell, H., 2018. Validation strategies for the interpretation of microstructure imaging using diffusion MRI. NeuroImage 182, 62–79.

Dyrby, T.B., Sogaard, L.V., Hall, M.G., Ptito, M., Alexander, D.C., 2013. Contrast and stability of the axon diameter index from microstructure imaging with diffusion mri. Magnetic Resonance in Medicine 70, 711–721.

D’Arceuil, H.E., Westmoreland, S., de Crespigny, A.J., 2007. An approach to high resolution diffusion tensor imaging in fixed primate brain. Neuroimage 35, 553–565.

Ferizi, U., Scherrer, B., Schneider, T., Alipoor, M., Eufracio, O., Fick, R., Deriche, R., Nilsson, M., Loya-Olivas, A., Rivera, M., Poot, D., Ramirez-Manzanares, A., Marroquin, J., Rokem, A., Potter, C., Dougherty, R., Sakaie, K., Wheeler-Kingshott, C., Warfield, S., Witzel, T., Wald, L., Raya, J., Alexander, D., 2017. Diffusion MRI microstructure models with in vivo human brain Connectom data: results from a multi-group comparison. NMR in Biomed 9.

Fieremans, E., H, J.J., A, H.J., 2011. White matter characterization with diffusional kurtosis imaging. Neuroimage 58, 177–188.

Hukkanen, V., Röyttä, M., 1987. Autolytic changes of human white matter: an electron microscopic and electrophoretic study. Experimental and molecular pathology 46, 31–39.

Innocenti, G.M., Vercelli, A., Caminiti, R., 2013. The diameter of cortical axons depends both on the area of origin and target. Cerebral Cortex 24, 2178–2188.

Jeffery, G., Erskine, L., 2005. Variations in the architecture and development of the vertebrate optic chiasm. Progress in retinal and eye research 24, 721–753.

Jenkinson, M., Bannister, P., Brady, M., Smith, S., 2002. Improved optimization for the robust and accurate linear registration and motion correction of brain images. Neuroimage 17, 825–841.

Jespersen, S.N., 2018. White matter biomarkers from diffusion mri. Journal of Magnetic Resonance 291, 127–140.

Jeurissen, B., Leemans, A., Tournier, J.D., Jones, D.K., Sijbers, J., 2013. Investigating the prevalence of complex fiber configurations in white matter tissue with diffusion magnetic resonance imaging. Human Brain Mapping 34, 2747–66.

Jeurissen, B., Tournier, J.D., Dhollander, T., Connelly, A., Sijbers, J., 2014. Multi-tissue constrained spherical deconvolution for improved analysis of multi-shell diffusion mri data. NeuroImage 103, 411–426.

Jones, D., Alexander, D., Bowtell, R., Cercignani, M., Dell’Acqua, F., McHugh, D., Miller, K., Palombo, M., Parker, G., Rudrapatna, U., et al., 2018. Microstructural imaging of the human brain with a ‘super-scanner’: 10 key advantages of ultra-strong gradients for diffusion mri. NeuroImage 182, 8–38.

Jones, D.K., Knösche, T.R., Turner, R., 2013. White matter integrity, fiber count, and other fallacies: the do’s and don’ts of diffusion mri. Neuroimage 73, 239–254.

Khan, A.R., Cornea, A., Leigland, L.A., Kohama, S.G., Jespersen, S.N., Kroenke, C.D., 2015. 3d structure tensor analysis of light microscopy data for validating diffusion mri. Neuroimage 111, 192–203.

Klawiter, E.C., Schmidt, R.E., Trinkaus, K., Liang, H.F., Budde, M.D., Naismith, R.T., Song, S.K., Cross, A.H., Benzinger, T.L., 2011. Radial diffusivity predicts demyelination in ex vivo multiple sclerosis spinal cords. Neuroimage 55, 1454–1460.

Lee, H.H., Yaros, K., Veraart, J., Pathan, J.L., Liang, F.X., Kim, S.G., Novikov, D.S., Fieremans, E., 2019. Along-axon diameter variation and axonal orientation dispersion revealed with 3d electron microscopy: implications for quantifying brain white matter microstructure with histology and diffusion mri. Brain Structure and Function.

Leergaard, T.B., White, N.S., De Crespigny, A., Bolstad, I., D’Arceuil, H., Bjaalie, J.G., Dale, A.M., 2010. Quantitative histological validation of diffusion mri fiber orientation distributions in the rat brain. PloS one 5, e8595.

Lefebvre, J., Delafontaine-Martel, P., Pouliot, P., Girouard, H., Descoteaux, M., Lesage, F., 2018. Fully automated dual-resolution serial optical coherence tomography aimed at diffusion mri validation in whole mouse brains. Neurophotonics 5, 045004.

Liu, M., Gross, D.W., Wheatley, B.M., Concha, L., Beaulieu, C., 2013. The acute phase of wallerian degeneration: longitudinal diffusion tensor imaging of the fornix following temporal lobe surgery. Neuroimage 74, 128–139.

Mitra, P.P., 1995. Multiple wave-vector extensions of the nmr pulsed-field-gradient spin-echo diffusion measurement. Physical Review B 51, 15074.

Novikov, D.S., Kiselev, V.G., Jespersen, S.N., 2018. On modeling. Magnetic resonance in medicine 79, 3172–3193.

O’Donnell, L.J., Westin, C.F., 2011. An introduction to diffusion tensor image analysis. Neurosurgery Clinics 22, 185–196.

Panagiotaki, E., Schneider, T., Siow, B., Hall, M.G., Lythgoe, M.F., Alexander, D.C., 2012. Compartment models of the diffusion mr signal in brain white matter: a taxonomy and comparison. Neuroimage 59, 2241–2254.

Paus, T., 2018. Imaging microstructure in the living human brain: a viewpoint. NeuroImage 182, 3–7.

Peters, A., 1970. The fixation of central nervous tissue and the analysis of electron micrographs of the neuropil, with special reference to the cerebral cortex, in: Contemporary Research Methods in Neuroanatomy. Springer, pp. 56–76.

Pichat, J., Iglesias, J.E., Yousry, T., Ourselin, S., Modat, M., 2018. A survey of methods for 3d histology reconstruction. Medical image analysis 46, 73–105.

Preibisch, S., Saalfeld, S., Tomancak, P., 2009. Globally optimal stitching of tiled 3d microscopic image acquisitions. Bioinformatics 25, 1463–1465.

Raffelt, D., Tournier, J.D., Rose, S., Ridgway, G.R., Henderson, R., Crozier, S., Salvado, O., Connelly, A., 2012. Apparent fibre density: A novel measure for the analysis of diffusion-weighted magnetic resonance images. NeuroImage 59, 3976–3994.

Riffert, T.W., Schreiber, J., Anwander, A., Knosche, T.R., 2014. Beyond fractional anisotropy: Extraction of bundle-specific structural metrics from crossing fiber models. NeuroImage 100, 176–191.

Salo, R.A., Belevich, I., Manninen, E., Jokitalo, E., Gröhn, O., Sierra, A., 2018. Quantification of anisotropy and orientation in 3d electron microscopy and diffusion tensor imaging in injured rat brain. NeuroImage 172, 404–414.

Scherrer, B., Schwartzman, A., Taquet, M., Sahin, M., Prabhu, S.P., Warfield, S.K., 2016. Characterizing brain tissue by assessment of the distribution of anisotropic microstructural environments in diffusion-compartment imaging (diamond). Magnetic resonance in medicine 76, 963–977.

Scherrer, B., Taquet, M., Schwartzman, A., St-Onge, E., Rensonnet, G., Prabhu, S.P., Warfield, S.K., 2017. Decoupling axial and radial tissue heterogeneity in diffusion compartment imaging, in: Proc. of the 25th Int Conf Inf Process Med Imaging (IPMI), pp. 440–452.

Schilling, K., Janve, V., Gao, Y., Stepniewska, I., Landman, B.A., Anderson, A.W., 2016. Comparison of 3d orientation distribution functions measured with confocal microscopy and diffusion mri. Neuroimage 129, 185–197.

Schindelin, J., Arganda-Carreras, I., Frise, E., Kaynig, V., Longair, M., Pietzsch, T., Preibisch, S., Rueden, C., Saalfeld, S., Schmid, B., et al., 2012. Fiji: an open-source platform for biological-image analysis. Nature methods 9, 676.

Schwartz, E.D., Duda, J., Shumsky, J.S., Cooper, E.T., Gee, J., 2005. Spinal cord diffusion tensor imaging and fiber tracking can identify white matter tract disruption and glial scar orientation following lateral funiculotomy. Journal of neurotrauma 22, 1388–1398.

Shemesh, N., Jespersen, S.N., Alexander, D.C., Cohen, Y., Drobnjak, I., Dyrby, T.B., Finsterbusch, J., Koch, M.A., Kuder, T., Laun, F., et al., 2016. Conventions and nomenclature for double diffusion encoding nmr and mri. Magnetic resonance in medicine 75, 82–87.

Sierra, A., Laitinen, T., Gröhn, O., Pitkänen, A., 2015. Diffusion tensor imaging of hippocampal network plasticity. Brain Structure and Function 220, 781–801.

Smith, R.E., Tournier, J.D., Calamante, F., Connelly, A., 2013. Sift: spherical-deconvolution informed filtering of tractograms. Neuroimage 67, 298–312.

Song, S.K., Sun, S.W., Ju, W.K., Lin, S.J., Cross, A.H., Neufeld, A.H., 2003. Diffusion tensor imaging detects and differentiates axon and myelin degeneration in mouse optic nerve after retinal ischemia. Neuroimage 20, 1714–22.

Sporns, O., 2011. The human connectome: a complex network. Annals of the New York Academy of Sciences 1224, 109–125.

Stanisz, G.J., Midha, R., Munro, C.A., Henkelman, R.M., 2001. Mr properties of rat sciatic nerve following trauma. Magnetic Resonance in Medicine: An Official Journal of the International Society for Magnetic Resonance in Medicine 45, 415–420.

Sun, S.W., Liang, H.F., Cross, A.H., Song, S.K., 2008. Evolving wallerian degeneration after transient retinal ischemia in mice characterized by diffusion tensor imaging. Neuroimage 40, 1–10.

Tournier, J.D., Calamante, F., Connelly, A., 2013. Determination of the appropriate b value and number of gradient directions for high-angular-resolution diffusion-weighted imaging. NMR in Biomedicine 26, 1775–1786.

Tournier, J.D., Mori, S., Leemans, A., 2011. Diffusion tensor imaging and beyond. Magnetic Resonance in Medicine 65, 1532–1556.

Tustison, N.J., Avants, B.B., Cook, P.A., Zheng, Y., Egan, A., Yushkevich, P.A., Gee, J.C., 2010. N4itk: improved n3 bias correction. IEEE transactions on medical imaging 29, 1310–1320.

Tyszka, J.M., Readhead, C., Bearer, E.L., Pautler, R.G., Jacobs, R.E., 2006. Statistical diffusion tensor histology reveals regional dysmyelination effects in the shiverer mouse mutant. Neuroimage 29, 1058–1065.

Veraart, J., Novikov, D.S., Christiaens, D., Ades-Aron, B., Sijbers, J., Fieremans, E., 2016. De-noising of diffusion mri using random matrix theory. NeuroImage 142, 394–406.

Wang, H., Zhu, J., Reuter, M., Vinke, L.N., Yendiki, A., Boas, D.A., Fischl, B., Akkin, T., 2014. Cross-validation of serial optical coherence scanning and diffusion tensor imaging: a study on neural fiber maps in human medulla oblongata. Neuroimage 100, 395–404.

Westin, C.F., Maier, S.E., Mamata, H., Jolesz, F.A., Kikinis, R., 2002. Processing and visualization for diffusion tensor MRI. Medical Image Analysis 6, 93–108.

Yang, G., Tian, Q., Leuze, C., Wintermark, M., McNab, J.A., 2018. Double diffusion encoding mri for the clinic. Magnetic resonance in medicine 80, 507–520.

Zaimi, A., Duval, T., Gasecka, A., Cote, D., Stikov, N., Cohen-Adad, J., 2016. Axonseg: open source software for axon and myelin segmentation and morphometric analysis. Frontiers in neuroinformatics 10, 37.

Zhang, H., Schneider, T., Wheeler-Kingshott, C.A., Alexander, D.C., 2012. Noddi: Practical in vivo neurite orientation dispersion and density imaging of the human brain. NeuroImage 61, 1000–1016.

